# T2T-CHM13 vs GRCh38: accurate identification of immunoglobulin isotypes from scRNA-seq requires a genome reference matched for ethnicity

**DOI:** 10.1101/2023.05.24.542206

**Authors:** Junli Nie, Julie Tellier, Ilariya Tarasova, Stephen L. Nutt, Gordon K. Smyth

**Affiliations:** Walter and Eliza Hall Institute of Medical Research, Parkville, Victoria 3052, Australia; Department of Medical Biology, University of Melbourne, Parkville, Victoria 3010, Australia; School of Mathematics and Statistics, University of Melbourne, Parkville, Victoria 3010, Australia

**Keywords:** Reference genome, T2T-CHM13, GRCh38, Immunoglobin genes, Clonal selection theory

## Abstract

Antibody production by B-cells is essential for protective immunity. The clonal selection theory posits that each mature B cell has a unique immunoglobulin receptor generated through random gene recombination and, when stimulated to differentiate into an antibody-secreting cell, has the capacity to produce only a single antibody specificity. It follows from this “one-cell-one-antibody” dogma, that single-cell RNA-seq profiling of antibody-secreting cells should find that each cell expresses only a single form of each of the immunoglobulin heavy and light chains. However, when using GRCh38 as the genome reference, we found that many antibody-secreting cells appeared to express multiple immunoglobulin isotypes. When the newly published T2T-CHM13 genome was used instead as the genome reference, every antibody-secreting cell was found to express a unique isotype, and read mapping quality was also improved. We show that the superior performance of T2T-CHM13 was due to its European origin matching the genetic background of the query samples. On the other hand, T2T-CHM13 failed to appropriately fit the “one-cell-one-antibody” dogma when applied to data derived from East Asia. Our results show that read assignment to human immunoglobulin isotype genes is very sensitive to the ancestral origin of the genome reference.

## Introduction

The human genome was first published in 2001(1–3) and has been continually improved over the past 20 years. For the past decade, the standard human reference genome in use by the UCSC Genome Browser, RefSeq, Ensembl and Gencode has been the GRCh38 genome released by the Genome Reference Consortium in 2013(4). Additional patches have been released regularly, most recently GRCh38.p14 in February 2022 (https://www.ncbi.nlm.nih.gov/datasets/genome/-GCF_000001405.40). GRCh38 has been accepted as a high quality representation of the 92% of the human genome that is euchromatic and accessible for gene transcription and has been the basis of most human genomic analyses over the past decade(3,4).

Recently, the telomere-to-telomere (T2T) consortium published the T2T-CHM13 assembly, the first essentially complete sequence of a human genome(5). The T2T-CHM13 assembly provides complete coverage of heterochromatic and repetitive regions of the human genome and has been shown to improve identification of SNPs and structural variations(6). Nevertheless, the differences between T2T-CHM13 and GRCh38 for protein-coding genes have been little explored. Most annotated genes and exons are covered by both T2T-CHM13 and GRCh38 and it is difficult to rigorously compare the accuracy of the alternative reference genomes for these common regions unless there is some sort of known ground truth against which to assess the comparison. In this article, we compare T2T-CHM13 and GRCh38 for an important gene family, immunoglobulin isotype genes, for which the “one cell, one antibody” dogma provides such ground truth.

B cells and immunoglobins generated by B cells form a major arm of the adaptive immune system(7). Antigens binding to B cell receptors, with suitable co-stimulatory signals, activate B cells to proliferate and differentiate into antibody-secreting cells (ASCs)(8). The antibodies produced by ASCs present a high binding affinity to their target antigens, neutralizing infectious threats, regulating the microbiota, or clearing dead cells from the body(9). Antibodies are encoded by the immunoglobulin genes (*IGH*, *IGK*, *IGL*)(10–12) and are expressed exclusively by the lymphocytes of the B cell lineage. An antibody molecule consists of two identical heavy chains and two identical light chains. Both heavy and light chains possess an N-terminal variable region and a C-terminal constant region, with the variable region determining antigen specificity. Based on the type of the constant region of the heavy chain, human immunoglobulins are divided into five major isotypes or classes, IgM, IgD, IgG, IgA, and IgE, among which IgG is further divided into IgG1, IgG2, IgG3, and IgG4 and IgA into IgA1 and IgA2 (Fig S1)(13).

An essential step in the formation of a functional humoral immune system is the somatic genomic recombination that integrates 2 or 3 of the segments from the immunoglobulin genes to a functional exon encoding the N-terminal variable region for a particular cell(14) (Fig S1). This process initiates in developing B cells in the bone marrow and occurs in an ordered fashion, where the *IGH* locus undergoes sequential Diversity (D) to Joining (J) recombination, followed by Variable (V) to DJ recombination, to produce a single VDJ exon encoding the N-terminal variable region of a heavy chain. Upon successful VDJ recombination in one chromosome, further recombination on the second chromosome is inhibited to ensure that the developing B cell only carries a single functional VDJ rearranged *IGH*, a process termed allelic exclusion(15,16). Following successful *IGH* rearrangement, developing B cells then undergo a similar (V-J) recombination process in either one of the two immunoglobulin light chain loci (*IGK*, *IGL*).

Successful VDJ-recombination results in the circulation of mature naïve B cells expressing two forms of immunoglobulin, IgM and IgD, generated by alternative splicing of the same VDJ exon to either an *IGHM* exon or an *IGHD* exon (Fig S1)(17). Once activated, B cells have the unique capacity to replace the IgM/IgD constant regions with other isotypes (IgG, IgA, or IgE) in a process called immunoglobulin class switch recombination (CSR). CSR results in the VDJ exon of the *IGH* being recombined to any one of the downstream exons encoding the constant regions of IgG, IgE or IgA by enzymatic deletion of all intervening sequences(18) (Fig S1). The sequence deletion physically ensures that only one isotype is expressed per cell. Key features of the system are that the antibody isotypes have distinct functions(19–24) and that CSR is differentially induced by infections(25–27), mRNA vaccines(28), autoimmune diseases(29) and anti-tumour responses(30). This process thus allows the type of antibody response to be tailored to meet the challenge at hand. At the completion of this stage, B cells undergo differentiation into antibody-secreting cells (ASCs), which transcribe large amounts of a shorter isoform of the *IGH* mRNA resulting in a secreted antibody form of the protein(31).

An important consequence of the VDJ-recombination process outlined above is that each B cell carries only a single functional and unique immunoglobulin molecule. This “one cell, one antibody” principle was captured in clonal selection theory proposed by Burnet (the 1960 Nobel laureate)(15), and first proven experimentally by Nossal(32), that underlies the modern industrial production of monoclonal antibodies for research and therapy. Together with the process of CSR, this theory implies that only one immunoglobulin isotype should be present within each single antibody-secreting plasma cell. Advances in single cell RNA sequencing methods now provide the opportunity to validate this classic immunological concept. Here we show that, when using the GRCh38 (referred to as hg38) as the genome reference for four public datasets, the results appear to violate the “one cell, one antibody” dogma. On the other hand, we found that the results agree with the “one cell, one antibody” theory perfectly if we switch the reference genome from hg38 to the newly published T2T-CHM13 (T2T) genome. Investigating this phenomenon further, we found that the sequence differences reflect the ethnic discrepancies between T2T and hg38. When using an ethnicity-matched genome reference, the identification and quantification of immunoglobulin genes are more accurate.

Given the essential roles of the immunoglobulin isotypes in human health, and the importance of mapping the functionality B cells in providing insights into the immune responses to infection, vaccination, allergy and in autoimmune conditions, it is necessary to identify them accurately in RNA sequencing data. These study highlights the importance of considering the reference genome in achieving this goal.

## Materials and Methods

### 10x Genomics Chromium single-cell datasets

10x-format FASTQ files were downloaded for each of five public single-cell datasets. The primary dataset focuses specifically on bone marrow plasma cells(33). Other datasets were derived from intestinal mucosae(34), tonsil(35), bone marrow(36), and intestinal mucosae from Chinese individuals(37).

The bone marrow plasma cell (BMPC) data was generated using Cell Ranger v5.0.0. Metadata is available from GEO accession GSM7181915 and FASTQ files from SRA accession SRX20001597.

Metadata for the intestine mucosae dataset is available from GEO accession GSM3972026 and FASTQ files were downloaded from SRA accession SRX6584269.

Metadata for the tonsil dataset is available from GEO accession GSM5051497 and a BAM file containing sequence reads was downloaded from SRA accession SRX9986858. The BAM file was converted to FASTQ files using the Cell Ranger v7.0.0 utility *bamtofastq*.

Metadata for the bone marrow dataset is available from GEO accession GSM3396184 and a BAM file was downloaded from SRA accession SRX4720060. The BAM file was converted to FASTQ files using the Cell Ranger v7.0.0 utility *bamtofastq*.

Metadata for the Chinese intestinal mucosae dataset is available from GEO accession GSM3433583 and GSM3433584, and FASTQ files were downloaded from SRA accessions SRX4896886 and SRX4896887.

### Cell Ranger reference genomes

To make a reference genome for Cell Ranger, a FASTA file of genomic sequences and a GTF file of gene annotation is required. For GRCh38, the FASTA file was GRCh38.primary_assembly.genome.fa and the backbone GTF file was gencode.v35.annotation.gtf, both downloaded from https://www.gencodegenes.org. For T2T, the FASTA file was chm13v2.0_maskedY.fa and the backbone annotation was UCSC GENCODEv35 CAT/Liftoff v2, downloaded from https://github.com/marbl/CHM13.

rtracklayer(38) was used to read the GTFs and to facilitate help with subsetting. Genes were matched between genomes by symbol. The analysis was restricted to protein-coding genes to make the results more reproducible between alternative annotations(39). Any gene not found in both GRCh38 and T2T annotations was also removed.

FASTA and GTF files for the two Chinese genomes were downloaded from https://ftp.ensembl.org/pub/rapid-release/species/Homo_sapiens/GCA_018471515.1/ and https://ftp.ensembl.org/pub/rapid-release/species/Homo_sapiens/GCA_018472605.1. The two genomes correspond to HG00438 and HG00621 in Figure 1h of Liao et al.(40), but are relabeled as Han-438 and Han-621 in this article to emphasize their Han Chinese ethnic origins. Gene annotation for the two Han genome references was restricted to protein-coding genes as for hg38 and T2T.

**Figure 1.**
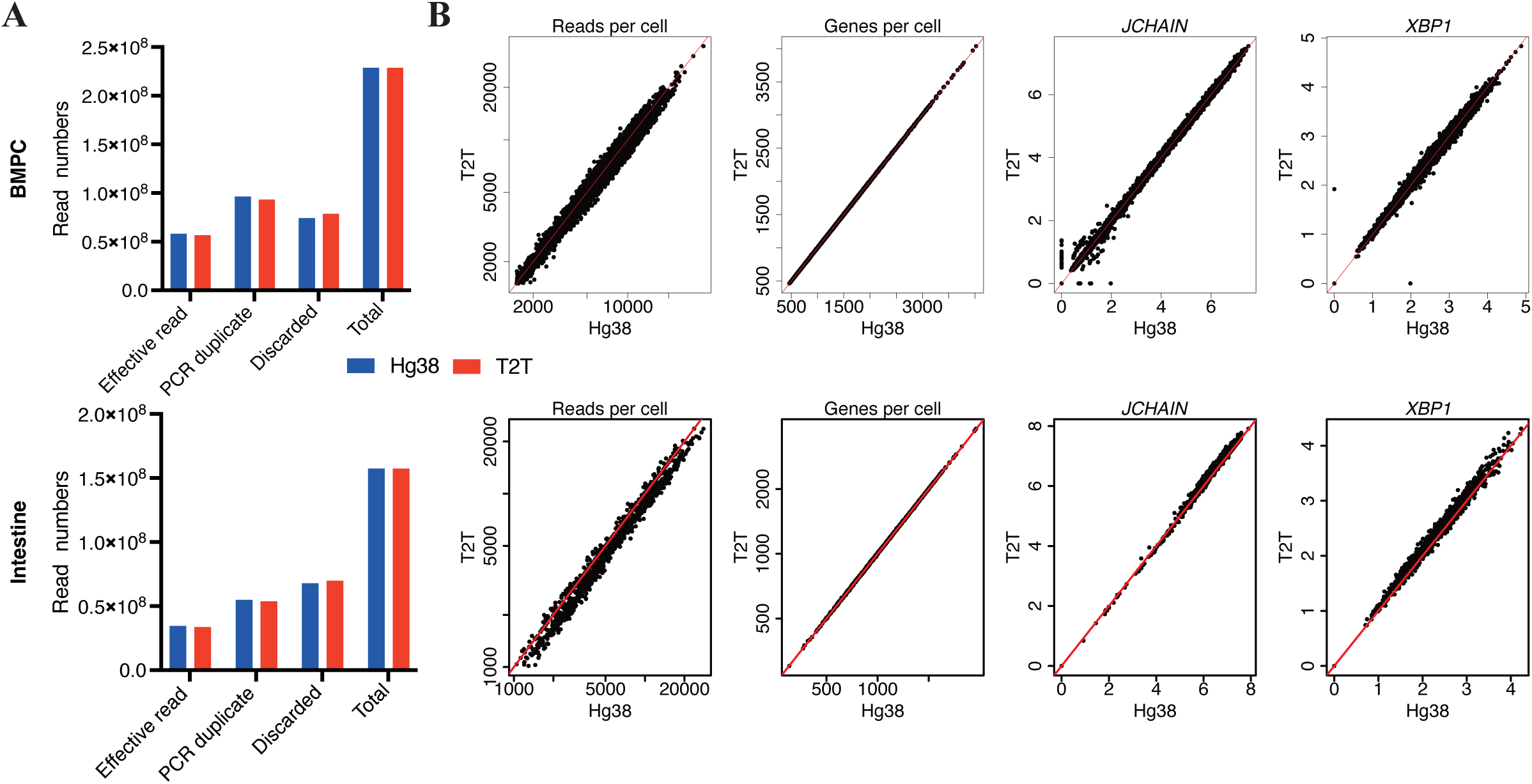
Mapping performance is comparable between hg38 and T2T. **A)** Classification of reads for the bone marrow plasma cell (BMPC, top) and intestinal mucosa (bottom) datasets for each reference genome. The barplots show the total read count for each sample and also the number of effective reads, the number of reads identified as PCR duplicates and the number of reads discarded for other reasons. Only effective reads were used for downstream analyses. **B)** Reads per cell, genes detected per cell, and expression of two plasma cell marker genes (*JCHAIN* and *XBP1*) for the BMPC (top) and intestinal mucosa (bottom) datasets. Here and subsequently, only bona fide ASCs are shown. The hg38 and T2T cell-wise counts and expression values are compared by scatter plots. Expression is represented as the log1p of reads per ten thousand (expression value = log1p(y/L*10^4^) where y is the read count for the gene and L is total read count for the cell.). Red lines in scatter plots represent the x=y line.

Finally, Cell Ranger references were made by *cellranger mkref* v7.0.0, with the FASTA files and filtered GTF files as inputs.

The custom T2T reference for Figure 6 was made by editing the GTF file in accordance with Figure 6A, the remaining process being the same.

### Cell Ranger mapping and quantification

FASTQ files are mapped to GRCh38 and T2T separately. Gene x barcode count matrices were generated by *cellranger count* v7.0.0.

To generate Figures 1A, 4A, 5A, 6B, S2, S10, S11D, S12A, S18 left, and S20A, BAM files generated by *cellranger count* were used to check the general mapping performance. The xf, GN, AS, and nM tags were extracted from the BAM files by samtools(41). For each read, GN indicates which gene the read is assigned to, xf indicates the alignment status (effective, PCR duplicate, or discarded), AS gives the mapping score, and nM the edit distance (https://www.10xgenomics.com/support/software/cell-ranger/latest/analysis/outputs/cr-outputs-bam, https://github.com/alexdobin/STAR/blob/master/doc/STARmanual.pdf).

### Quality control and dimension reduction plots

The matrix output from *cellranger count* were further processed following the general Seurat(42) pipeline, in which quality control, normalisation, and dimension reduction is performed. Quality control involved 4 steps, (1) cells that have >10% mitochondrial gene expression were removed; (2) ribosomal genes were removed; (3) genes that are expressed in fewer than 0.5% (0.25% for the intestine mucosae dataset) of total cells were removed; (4) and cells with too few reads (usually 1500 as threshold) or expressed genes (usually 450 as threshold) were removed (Figure S3, and Table S1). Finally, only cells that have >10% immunoglobulin gene expression (Figure S4) and appear in both hg38 and T2T output are defined as antibody-secreting cells for downstream analysis and for generating Figures 1B, 2, 3A, 4B, and 5BC (scatter plots) together with Figures S5, S6, S7, S8, S9, S11A, S12B, S14B, S15B, S16B, S17B, and S18 scatter plots.

To generate Figure 6CDE, and Figure 7BC together with Figures S20BC and S21, the similar process was performed but with the custom references.

### Alignment track plot visualisation

To generate Figures 3B, S11B, 4CD, and 5C, BAM files of particular genes and cells were generated by *samtools* based on the read names, CR tag (representing cell barcodes), GN tag, and xf tag (only effective reads and PCR duplicate reads were included). Alignment track plots were produced from the BAM files with the R package Gviz(43).

### Sequence alignment

To determine the allotype alleles, an online tool, IMGT/BlastSearch(44), was used based on the databases of IMGT(45,46).

A Plasmid Editor (ApE) software(47) (https://jorgensen.biology.utah.edu/wayned/ape) was used to generate Figures 3C, S11C, 4EF, S13, S14A, S15A, S16A, S17A, and S19.

## Results

### Mapping performance is comparable between hg38 and T2T

To compare the performance of hg38 and T2T in isotype identification, two human 10x Genomics Chromium 3’ scRNA-seq datasets were selected, one generated in-house from enriched bone marrow plasma cells (BMPCs, from one individual (ref 33) and the other from intestinal mucosae of one individual published in Martin et al(34). Two supplementary 10x Genomics Chromium 3’ scRNA-seq datasets from King et al(35) (one tonsil samples) and Oetjen et al(36) (one bone marrow sample, BM) were also interrogated to demonstrate the reproducibility of the results. Equivalent custom Cell Ranger references were constructed from the hg38 and T2T genomes, then cell-by-gene matrices of read counts were obtained from Cell Ranger. Examining the alignment output BAM files, mapping performances of hg38 and T2T at the read level were similar, with approximately the same numbers of effectively mapped sequence reads and PCR duplicates (Fig 1A and Fig S2A). The edit distances and mapping scores of the reads were also comparable (Fig S2B). After the read alignment and cell demultiplexing, quality control (QC) was performed on all single cells (Table S1, Fig S3). The QC metrics were similar between the hg38 and T2T output (Fig S3). These results show that there are no global differences between the hg38 and T2T genomes and that the two genomes have been processed in equivalent ways in our pipeline. After QC, bona fide antibody-secreting cells (ASCs) were selected based on immunoglobulin gene expression and only cells appearing in both hg38 and T2T outputs were retained (Fig S4). Read and gene coverage per ASC cell was also comparable between hg38 and T2T (Fig 1B and Fig S5). In this study, ASCs are the target cells for isotype detection. Expression levels of two key marker genes, *JCHAIN*(48), and *XBP1*(49) were similar between the hg38 and T2T outputs in all datasets (Fig 1C and Fig S5B). Taken together, the general qualities of single ASCs have few differences between the hg38 and T2T reference genomes.

### T2T but not hg38 identifies immunoglobulin isotypes correctly

We next examined expression levels of the Immunoglobin Heavy Chain (*IGH*) genes that are associated with isotype switching of ASCs. Plotting the expression of *IGHA*, *IGHD*, *IGHE*, *IGHG*, and *IGHM* in the ASCs did not reveal any unexpected patterns (Fig S6). Almost every ASC can be seen to express exactly one of these variants, and the small number of exceptions can be interpreted as likely doublet cells (Fig S6). The pattern was similar for both hg38 and T2T. The proportion of reads assigned to IGHM, IGHD and IGHE for each ASC correlated closely between hg38 and T2T (Fig S7).

Drilling down to the *IGH* subclass genes showed a different story, however. UMAP plots of ASC cells in the BMPC dataset, overlaid with *IGHG1*, *IGHG2*, *IGHG3*, and *IGHG4* expression, showed that most cells expressed exactly one isotype gene when T2T was the reference genome, but many cells unexpectedly expressed two isotypes simultaneously with hg38 as the reference (Fig 2A). A scatterplot of *IGHG1* vs *IGHG2* shows the problem very clearly (Fig 3A). With T2T as the reference, a handful of doublet cells can be seen, but almost all cells express exactly one isotype and are positioned on the vertical or horizontal axes. With hg38 as the reference, a large number of cells (759 cells) are positioned on a non-vertical trend line and apparently express both *IGHG1* and *IGHG2* (Fig 3A left). The double-positive cells cannot be interpreted as doublet cells because the frequency is so much greater than that observed for doublet cells in Fig S6, and because doublet cells would be randomly positioned rather than systematically arranged on a trend line. The same problem occurs with double-positive cells for *IGHG1* and *IGHG3* (Fig S8, Fig S11A). The hg38 output generates 1141 *IGHG1^+^IGHG3^+^* double-positive cells, on a broad diagonal trend line, whereas the T2T output shows only single-positive cells apart from a very small number of apparent doublets (Fig S11A).

**Figure 2.**
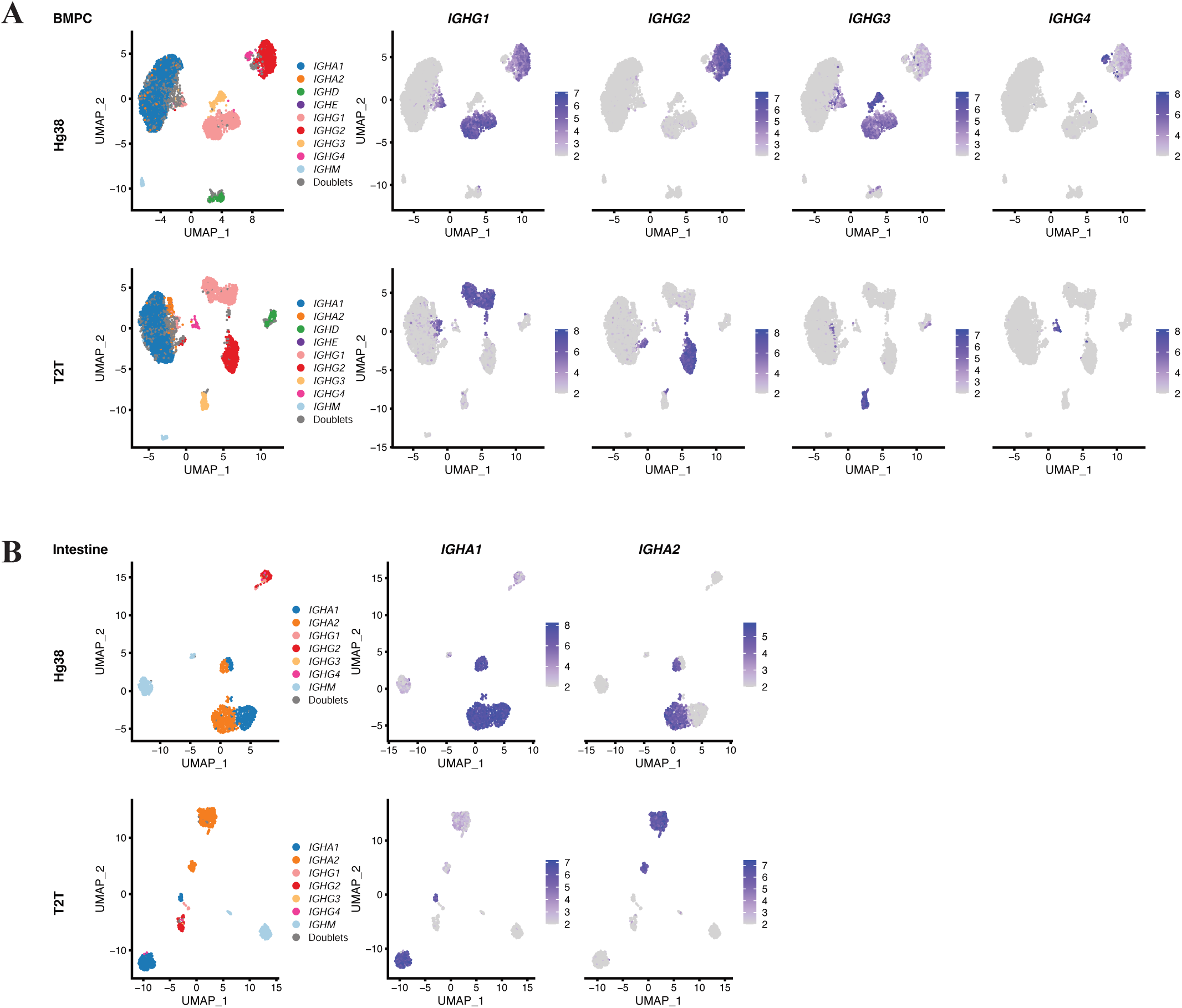
The T2T genome identifies immunoglobulin isotypes more accurately. **A)** UMAP dimension reduction plots of individual ASCs for the BMPC dataset. The left plots show the isotype assigned to each cell. The isotype assignment was based on the T2T output, in which if an isotype accounts for >85% expression of all isotype gene expression in a single cell, that isotype will be assigned to this single cell. In the right 4 columns, cells are coloured by expression level of the indicated gene for a particular IgG subclass. Plots are shown for hg38 (top) and T2T (bottom). Here expression values are represented as the log1p of reads per ten thousand (expression value = log1p(y/L*10^4^) where y is the read count for the gene and L is total read count for the cell.) **B)** UMAP dimension reduction plots of individual ASCs for the intestinal mucosa dataset. The left plots show the isotype assigned to each cell. The isotype assignment was based on the T2T output, in which if an isotype accounts for >85% expression of all isotype gene expression in a single cell, that isotype will be assigned to this single cell. In the right 2 columns, cells are coloured by expression level of the indicated gene for a particular IgA subclass. Plots are shown for hg38 (top) and T2T (bottom). Here expression values are represented the same as (A).

A similar situation occurs for the IgA isotype genes, *IGHA1* and *IGHA2*. Plotting *IGHA1* vs *IGHA2* expression for ASCs in the intestine mucosa dataset showed that half the cells were double-positive for hg38 but almost none for T2T (Fig 2B, 5C). The same pattern is seen for all the datasets (Fig S9).

The high rate of double-positive cells in the hg38 output is not compatible with the “one cell, one antibody” theory, and indicates incorrect alignment of sequence reads to the hg38 genome.

### Double-positive cells in the hg38 output should be single-positive

To explore why the T2T genome performs better for isotype identification, we focused first on *IGHG1^+^IGHG2^+^* double-positive cells and examined track graphics for the corresponding read alignments. The cells highlighted in red in Figure 3A show high expression of both *IGHG1* and *IGHG2* in the hg38 output but express only IGHG2 in the T2T output. To understand why this occurs, we examined the alignment positions of all the relevant reads in the hg38 output. Due to the Chromium 3’ sequencing, most of the reads from immunoglobulin transcripts were mapped to constant regions encoding the isotype genes (Fig S1). Reads were in particular mapped to the most 3’ exon of the secreted isoforms, which for *IGH* is the CH3-CHS exon (Constant Heavy chain exon 3). The left panel of Figure 3B shows read alignment coverage plots for the CH3-CHS exons of *IGHG1* and *IGHG2* in the hg38 output. The plots show all reads from the “red” cells in Figure 3A that map to either *IGHG1* or *IGHG2* in the hg38 output. The reads mapping to *IGHG1* are responsible for *IGHG1* expression in cells that are primarily *IGHG2* in the hg38 output.

**Figure 3.**
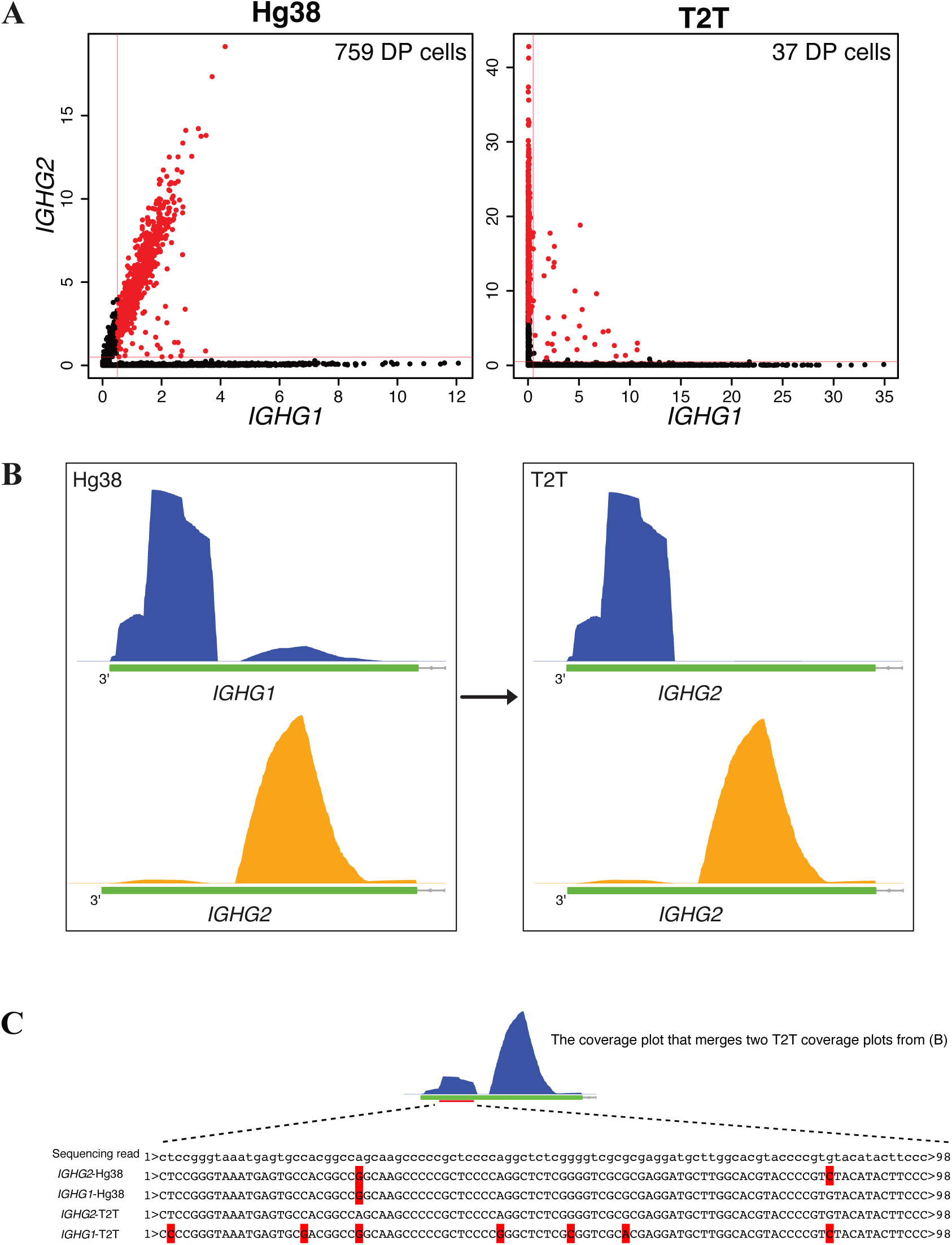
Separating *IGHG1^+^* and *IGHG2^+^* plasma cells. **A)** Scatter plots correlating *IGHG1* and *IGHG2* expression for BMPCs. Left plot for hg38 and right plot for T2T. The *IGHG1*+*IGHG2*+ cells of the hg38 output are highlighted in both plots. The horizontal and vertical red lines represent the thresholds for positive cells of an isotype gene. Here, the thresholds are 0.5% for both *IGHG1* and *IGHG2*. The numbers of double-positive (DP) cells for both outputs are also shown. **B)** Coverage plots for reads from the highlighted cells in (A). Top left plot shows reads mapped to *IGHG1* using hg38 and bottom left shows reads mapped to *IGHG2* using hg38. Only the CH3-CHS exon of the genes are shown. The plots on the right show coverage plots for the same reads but using T2T. The top right plot shows that all the reads mapped to *IGHG1* using hg38 map instead to *IGHG2* using T2T. The bottom right plot shows that all reads mapped to *IGHG2* using hg38 map also map to *IGHG2* using T2T. **C)** Alignment results for one illustrative sequence read incorrectly assigned to *IGHG1* by hg38 in (B). The full sequence of the read is shown together with the best matches to the *IGHG2*-hg38, *IGHG1*-hg38, *IGHG2*-T2T and *IGHG1*-T2T genome regions. Mismatched bases are marked in red. The read maps perfectly to *IGHG2*-T2T and, using T2T, the assignment to *IGHG2* is unambiguous. Using hg38, however, the result is reversed with fewer mismatches for *IGHG1*-hg38 than for *IGHG2*-hg38. The coverage plot merges the right two plots from (B) on the same scale, i.e., shows *IGHG2*-T2T coverage for all reads mapped by hg38 to either *IGHG1* or *IGHG2*. The aligned position of the illustrative read is shown by a red bar.

The right panel of Figure 3B shows the T2T coverage for the same reads as in the left panel. When aligned to T2T, the reads in the main *IGHG1* peak in the hg38 output were remapped to the homologous region of *IGHG2* instead of to *IGHG1*. The edit distances and mapping scores of these reads showed fewer mismatches and higher mapping quality when realigned to T2T instead of hg38 (Fig S10).

Comparing the consensus sequence of reads around the 3’ end region with the sequence of the 3’ end regions of *IGHG1* and *IGHG2* from both hg38 and T2T, the numbers of mismatches (red bases) were 2, 1, 0, 7, for *IGHG2*-hg38, *IGHG1*-hg38, *IGHG2*-T2T, and *IGHG1*-T2T, respectively (Fig 3C). Within hg38 output, *IGHG1* had fewer mismatches than *IGHG2*, thus those reads were mapped to *IGHG1* causing the double-positive cells, however, in T2T output, it was the *IGHG2* that had 0 mismatch, hence reads were mapped to *IGHG2* (Fig 3C). Taking these together, the *IGHG1*^+^*IGHG2*^+^ cells in the hg38 output should be *IGHG2* single-positive cells, because the reads in the 3’ end region from *IGHG2* single-positive cells in the real world are mapped to *IGHG1* incorrectly when using hg38 as genome reference.

Using a similar checking process, it is apparent that the *IGHG1*^+^*IGHG3^+^* double-positive cells, in the hg38 output, should be *IGHG1* single-positive cells (Fig S11).

### The T2T reference results in more IGHG reads than hg38

In addition to higher accuracy in isotype identification, the T2T output also has more reads that are mapped to IGHG genes. Checking the BAM files, there were more effective reads and PCR duplicates mapped to IGHG genes in the T2T output than the hg38 output because considerable numbers of reads were discarded in the hg38 output (Fig 4A and Fig S12A). At single cell level, many cells in the T2T output gained more IGHG gene expression than hg38 (Fig 4B and Fig S12B). Rechecking the *IGHG1*^+^ (red cells) of T2T output indicated in Fig S11A, in addition to 501294 reads that were also mapped to *IGHG1* or *IGHG3* in the hg38 output (Fig S11B, the right two track plots), 1768443 extra reads around the *IGHG1* stop codon were recovered (Fig 4C). Similarly, for the *IGHG2*^+^ (red cells) of T2T output indicated in Fig 3, in addition to 455966 reads that were also mapped to *IGHG2* or *IGHG1* in hg38 output (Fig 3B, the right two track plots), 1868398 extra reads around *IGHG2* stop codon region were recovered (Fig 4D). Tracking those extra recovered reads revealed that they were discarded in the hg38 output because they mapped to homologous regions of *IGHG1* and *IGHG2* and, hence, were discarded by the Cell Ranger software as multi-mapping reads with no unique best location. In contrast, the sequences of T2T, at the same locations, were distinguishable between *IGHG1* and *IGHG2* so reads could be mapped to *IGHG1* (Fig 4E) or *IGHG2* (Fig 4F) without ambiguity.

**Figure 4.**
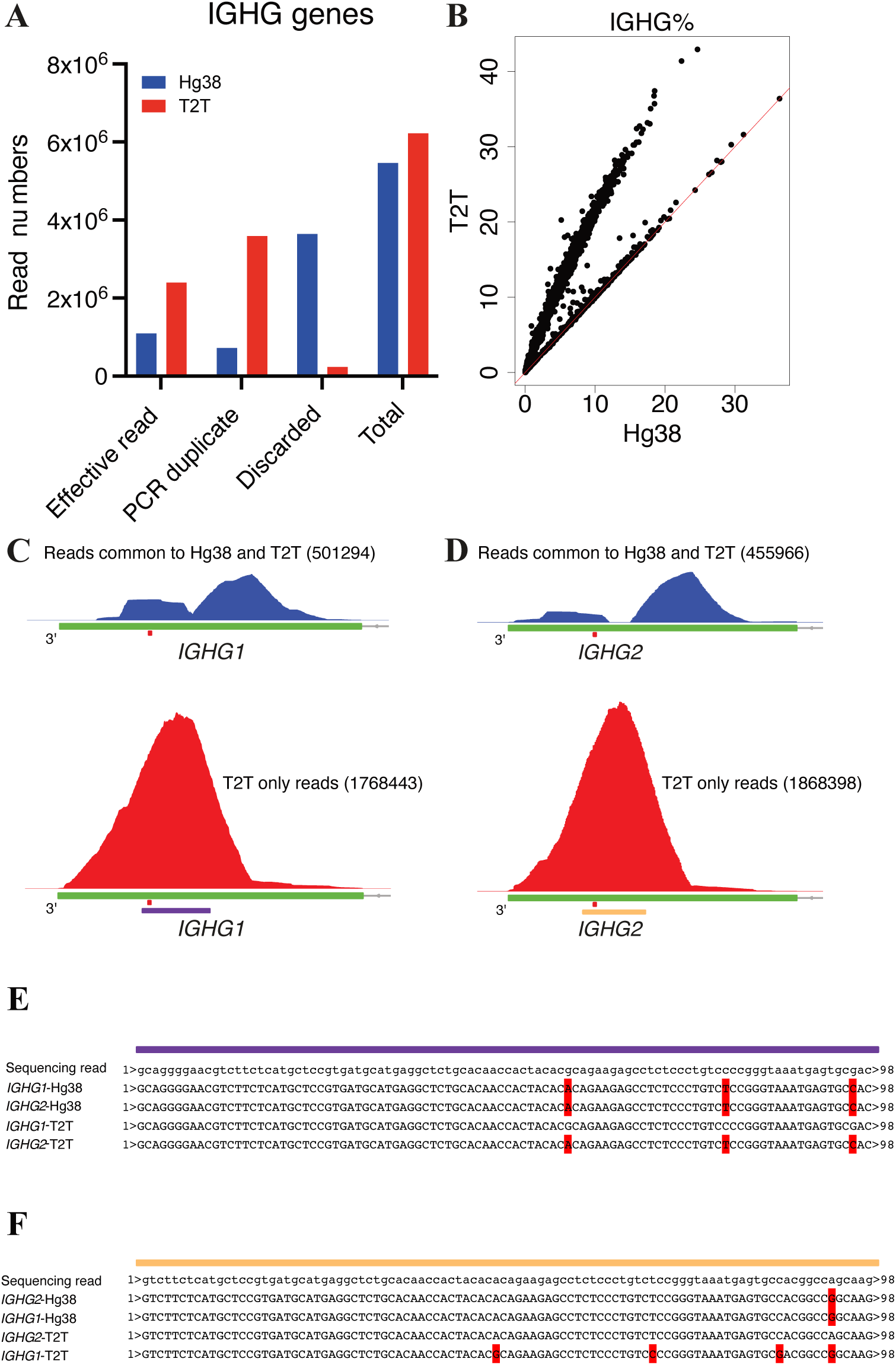
T2T reference results in more reads that are mapped to IGHG genes than hg38. **A)** Classification of reads that are assigned to IGHG genes for the BMPC dataset for each reference genome. The barplots show the total read count of IGHG genes and also the number of effective reads, the number of reads identified as PCR duplicates and the number of reads discarded for other reasons. **B)** Percentage of IGHG gene reads per cell for the BMPC dataset. Hg38 and T2T results are compared by scatter plots. Red lines in scatter plots represent the x=y line. **C)** Top, the coverage plot from Fig S11C with the same scale as the bottom coverage plot; bottom, the coverage plot of reads that map to *IGHG1* perfectly using T2T but with ambiguity when using hg38. The small red bar denotes the location of the stop codon. The numbers of reads that are inputted to generate the indicated coverage plots are also shown for the aim of comparison. The purple bar indicates the location of the sequencing read of (E). **D)** Top, the coverage plot from Fig 3C with the same scale as the bottom track plot; bottom, the coverage plot of reads that map to *IGHG2* perfectly using T2T but with ambiguity when using hg38. The small red bar denotes the location of the stop codon. The numbers of reads that are inputted to generate the indicated coverage plots are also shown for the aim of comparison. The yellow bar indicates the location of the sequencing read of (F). **E)** Alignment results for one illustrative sequence read with assignment ambiguity between *IGHG1* and *IGHG2* when using hg38 in (C). The full sequence of the read is shown together with the best matches to the *IGHG1*-hg38, *IGHG2*-hg38, *IGHG1*-T2T and *IGHG2*-T2T genome regions. Mismatched bases are marked in red. The read maps perfectly to *IGHG1*-T2T and, using T2T, the assignment to *IGHG1* is unambiguous. Using hg38, however, the result is ambiguous with the same number of mismatches between *IGHG1*-hg38 and *IGHG2*-hg38. **F)** Alignment results for one illustrative sequence read with assignment ambiguity between *IGHG1* and *IGHG2* when using hg38 in (D). The full sequence of the read is shown together with the best matches to the *IGHG2*-hg38, *IGHG1*-hg38, *IGHG2*-T2T and *IGHG1*-T2T genome regions. Mismatched bases are marked in red. The read maps perfectly to *IGHG2*-T2T and, using T2T, the assignment to *IGHG2* is unambiguous. Using hg38, however, the result is ambiguous with the same number of mismatches between *IGHG2*-hg38 and *IGHG1*-hg38.

To further understand the discrepancies of IGHG genes between T2T and hg38, the sequences of the CH3-CHS exons of *IGHG3*, *IGHG1*, *IGHG2*, and *IGHG4* from both T2T and hg38 were compared. There were in total 34 variants in the CH3-CHS exons among 4 IGHG genes in T2T (Fig S13A) but 28 variants in hg38 (Fig S13B), therefore the sequences of T2T were more distinguishable than hg38.

For *IGHG3*, the sequences of *IGHG3* of T2T and hg38 were nearly identical except the only one variant near the end of the 3-UTR (Fig S14A), hence, the expression levels of *IGHG3* in bone fide *IGHG3*+ cells (the dots around the x=y red line in Fig S14B) were comparable between T2T and hg38 (Fig S14B). The dots outside the x=y lines of Fig S14B represented the *IGHG1*+ cells, part of whose reads were assigned to *IGHG3*-hg38.

For *IGHG1*, the sequences were different between T2T and hg38 for both protein coding regions (5 mismatches) and 3-UTRs (8 mismatches) (Fig S15A), therefore the alignments and gene assignments of reads changed as shown in Fig 3C (*IGHG1*-hg38 to *IGHG2-T2T*), Fig S11C (*IGHG3*-hg38 to *IGHG1*-T2T) and Fig 4E (extra reads for *IGHG1*-T2T), and the consequent expression levels were also altered between T2T and hg38, T2T output having higher expression in *IGHG1*+ cells (Fig S15B).

For *IGHG2*, there was only one variant in the protein coding regions but 3 variants in the 3-UTRs (Fig S16A). The number of mismatches between hg38 and T2T were fewer than *IGHG1* but also resulting in that T2T had higher *IGHG2* expression in *IGHG2* single-positive cells (Fig S16B).

For *IGHG4*, T2T has two copies, which is found in 44% of humans(50). For the copy 1 (the 5’ upstream one), compared with hg38, it had two variants in the protein coding region but shared the same 3-UTR sequence (Fig S17A). For the copy 2, it had a nearly identical protein coding region as hg38 except one variant but a distinct 3-UTR (containing 6 variants) (Fig S17A). Thus, most of reads that could be mapped to *IGHG4* of hg38 could also be assigned to either one of *IGHG4* genes of T2T, which was proved by the comparable expression levels of *IGHG4* between T2T and hg38 (Fig S17B).

To conclude, the sequence differences of *IGHG1* and *IGHG2* between hg38 and T2T brought about the fewer multi-mapping reads and higher IGHG gene expression in the T2T output.

### T2T yields fewer IGHA reads but better distinguishes *IGHA1* and *IGHA2*

Unlike IGHG, there were fewer reads mapped to IGHA genes in the T2T output compared with hg38 (Fig 5A, 5B left, Fig S18 left two columns). This read reduction was associated with lower *IGHA1* expression (Fig 5B middle, Fig S18 third column) but higher *IGHA2* expression (Fig 5B right, Fig S18 fourth column).

**Figure 5.**
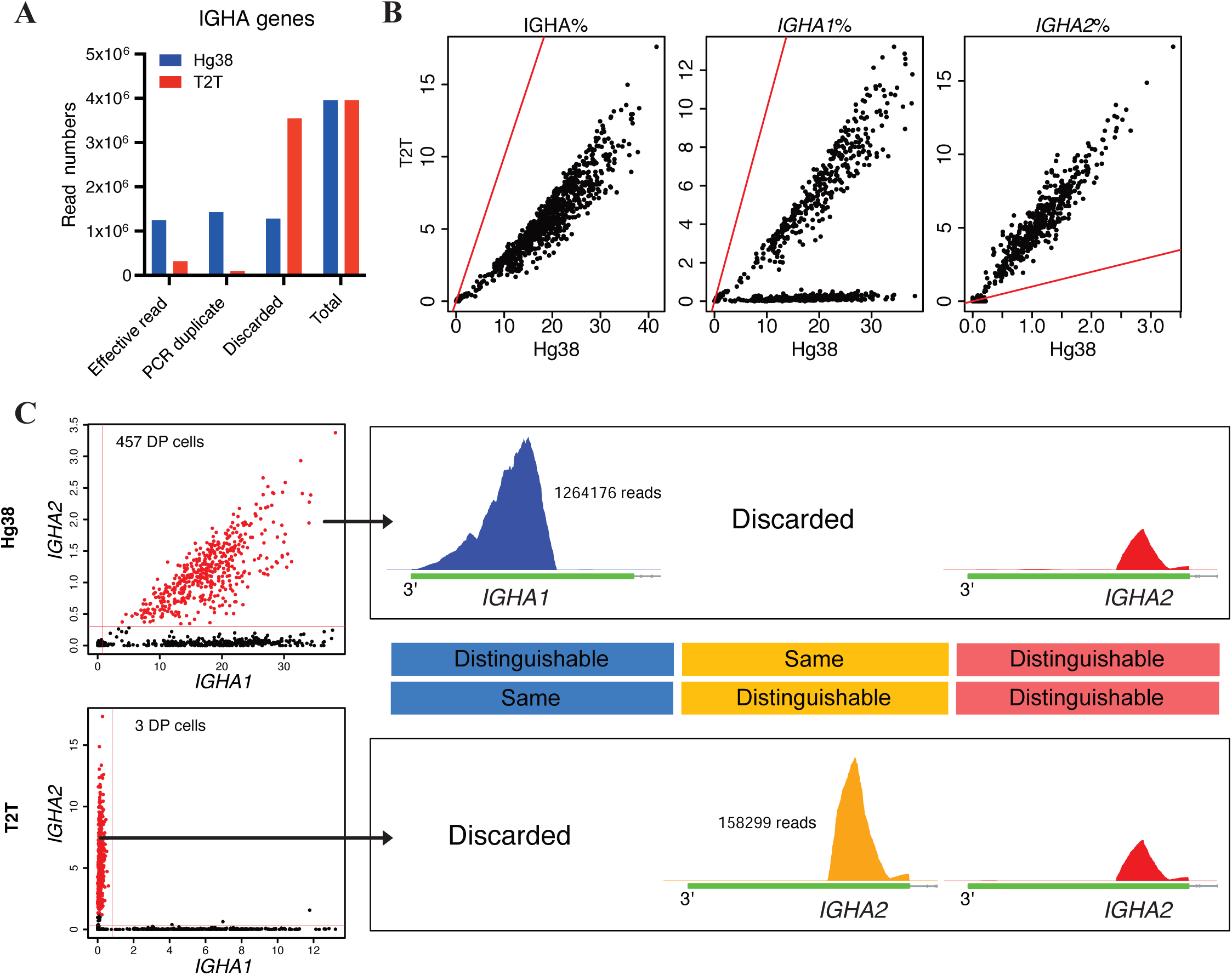
T2T output has fewer IGHA reads but better distinguishes *IGHA1* from *IGHA2*. **A)** Classification of reads that are assigned to IGHA genes for the intestinal mucosa (intestine) dataset for each reference genome. The barplots show the total read count of IGHA genes and also the number of effective reads, the number of reads identified as PCR duplicates and the number of reads discarded for other reasons. **B)** Percentage of IGHA gene reads per cell, *IGHA1* reads per cell, and *IGHA2* reads per cell for the intestine dataset. Hg38 and T2T results are compared by scatter. Red lines in scatter plots represent the x=y line. **C)** The left scatter plots correlating expression levels of *IGHA1* and *IGHA2* using hg38 (top) and T2T (bottom). The *IGHA1*+*IGHA2*+ cells of the hg38 output are highlighted in both plots. The horizontal and vertical red lines represent the thresholds for positive cells of an isotype gene. Here, the thresholds are 0.8% and 0.3%, respectively for *IGHA1* and *IGHA2*. The numbers of double-positive (DP) cells for both outputs are also shown. In the right, the plots on the top show coverage plots for reads aligned using hg38 from the highlighted cells in the scatter plots. Top left plot shows reads mapped to *IGHA1* using hg38 (these reads are discarded when using T2T because of mapping ambiguity) and top right shows reads mapped to *IGHA2* using hg38 (these reads also map *IGHA2* using T2T). Only the CH3-CHS exon of the genes is shown. The plots on the bottom show coverage plots for reads aligned using T2T. The bottom middle plot shows reads that map *IGHA2* perfectly, and using T2T, the assignment to *IGHA2* is unambiguous (these reads are discarded when using hg38 due to mapping ambiguity). The bottom right plot shows that all reads mapped to *IGHA2* using hg38 map also map to *IGHA2* using T2T. The numbers of reads that are inputted to generate the indicated coverage plots are also shown. A schematic plot of the CH3-CHS exons of *IGHA1* or *IGHA2* is between the hg38 and T2T coverage plots. The text explains the result of sequence alignment between *IGHA1* and *IGHA2* for the T2T or hg38 genomes. The colours used here have the same meaning as the coverage plots, i.e., the reads of coverage plots with one colour locate on the region in the same colour.

To understand this phenomenon in detail, we aligned the sequences of the CH3-CHS exons of *IGHA1* and *IGHA2* in the hg38 and T2T genomes, and divided them into 3 regions, marked as “blue”, “orange”, and “red” respectively in Figures 5C and S19.

In the “blue” region, the *IGHA1* and *IGHA2* sequences are identical in the T2T genome but slightly different in hg38. Reads mapping to this region were therefore discarded in the T2T output, because of multi-mapping, but mapped to *IGHA1* in the hg38 output (Fig 5C). The “blue” region includes the 3-UTRs of the two genes, together with the 3’ tail of the coding-regions where there are 9 variants between *IGHA1* and *IGHA2* in hg38 but none in T2T (Fig S19).

In the “orange” region, the *IGHA1* and *IGHA2* sequences are identical in the hg38 genome but distinguishable in T2T, so reads mapping this region are discarded in the hg38 output but mapped to *IGHA2* in the T2T output (Fig 5C, Fig 5B right). There is one variant distinguishing *IGHA1* from *IGHA2* in T2T (Fig S19), defining the orange region (Fig 5C and Fig S19).

Sequences in the “red” region are distinguishable in both hg38 and T2T, so reads mapping to this region were mapped to the appropriate genes correctly in both outputs (Fig 5C).

Taking these observations together, we can see that true *IGHA2* cells have become false *IGHA1^+^IGHA2^+^* double-positive cells in the hg38 output, because reads from those cells mapping to the “blue” region were incorrectly assigned to *IGHA1*. The same reads were discarded in the T2T output, resulting in lower *IGHA1* expression and avoidance of false double-positive cells. Meanwhile, T2T sequences are distinguishable in the “orange” region, resulting in higher *IGHA2* expression.

### A custom annotation reference for T2T can rescue multi-mapped reads mapped to IGHA genes

To quantify IGHA genes expression more precisely and rescue reads mapped to the “blue” region (Fig 5C), a custom annotation reference for T2T was created, in which the CH3-CHS exons of *IGHA2* and *IGHA1* were split into two “exons”, respectively (Fig 6A). The 3’ ones of the newly constructed exons, sharing the identical sequences (Fig S19), were named as *IGHA*, a new artificial gene, and were approximately equivalent to the “blue region” (Fig S19). When reads were mapped to *IGHA* exons, they would not be discarded even though they were multi-mapped reads because Cell Ranger retrieved multi-mapped reads if the regions they were mapped to pointed to the same gene, in this case, it is the *IGHA*. After this custom editing, effective reads assigned to IGHA genes were largely increased without influencing the general effective read numbers (Fig 6B, Fig S20A, and S20B). At the single cell level, the expression of general IGHA genes also increased (Fig 6C, the left, and Fig S20B), but the expression of both *IGHA1* and *IGHA2* decreased (Fig 6C, the middle and the right, and Fig S20B), implying that the increase in general expression was attributed to the new *IGHA* gene. Looking at the scatter plots of the expression levels of *IGHA1*, *IGHA2*, and *IGHA*, most of the *IGHA1*+ or *IGHA2*+ single cells are also *IGHA* positive, indicating that those single-positive had gained the rescued reads from the *IGHA* exons (Fig 6D and Fig S20C). Inheriting the advantages of T2T, this custom reference did not produce *IGHA1* and *IGHA2* double-positive cells (Fig 6D and Fig S20C) and had a well-separated clustering based on the isotype expression (Fig 6E). Therefore, the custom annotation reference could effectively overcome the issue of lost IGHA gene reads.

**Figure 6.**
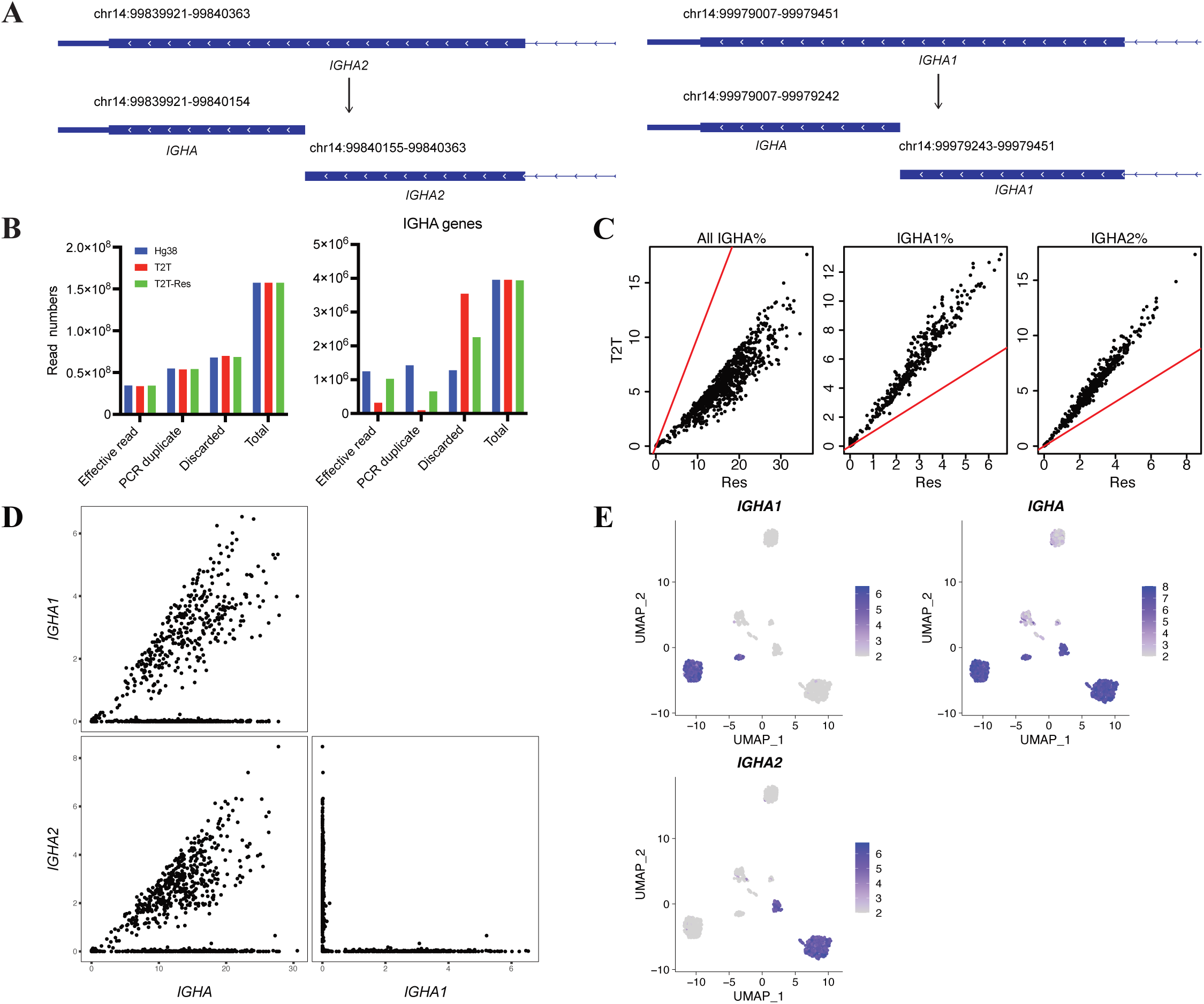
A custom annotation reference for T2T can rescue multi-mapped reads mapped to IGHA genes. **A)** Schematic plots showing the CH3-CHS exons of *IGHA2* and *IGHA1*, and how they are split into two exons, respectively. Coordinates of the exons are shown in the top of each exon. Gene names of the exons are shown in the bottom of each exon. Within each exon, the arrows denote the direction of the gene, the thicker blue bar indicate the protein coding region, and the thinner blue bar indicate the 3-UTR region. **B)** Classification of reads that are of the whole dataset (left) or are assigned to IGHA genes (right) for the intestinal mucosa datasets for each reference. The barplots show the total read count and also the number of effective reads, the number of reads identified as PCR duplicates and the number of reads discarded for other reasons. Only effective reads were used for downstream analyses. **C)** Percentage of IGHA gene reads (left), *IGHA1* reads (middle), and *IGHA2* reads (right) per cell for the intestine dataset. Hg38 and T2T results are compared by scatter. Red lines in scatter plots represent the x=y line. T2T-res is shown as “res” in plots. **D)** Paired scatter plots correlating the expression levels of pairs of IgA subclass genes for intestinal mucosa ASCs. Plots are shown for the output using the custom reference based on T2T. Here expression is represented as reads per hundred (RPH). **E)** UMAP dimension reduction plots of individual ASCs for the intestinal mucosa dataset. Cells are coloured by expression level of the indicated gene for a particular IgA subclass. Plots are shown for the output using the custom output based on T2T. Here expression values are represented the same as Fig 1C.

### The impact of allele differences on isotype calling

The above results demonstrate that sequence discrepancies caused different alignment outcome between hg38 and T2T for reads mapped to isotype genes. To determine the origins of those sequence variants, the full sequences including protein-coding regions and 3-UTRs of each IGHG and IGHA gene from both hg38 and T2T were aligned to the database of IMGT (https://www.imgt.org/blast/). The alignments indicated that hg38 and T2T represented different alleles for each IGHG and IGHA gene (Fig 7A). Bashirova et al.(51) has performed a comprehensive analysis of allele distribution of 3 IGHG genes, *IGHG1*, *IGHG2*, and *IGHG3*, between different ethnicities and showed that the most common haplotype of these 3 genes for European American was *IGHG3*\*11/*IGHG1*\*03/*IGHG2*\*02, the same haplotype as T2T, while the hg38 haplotype *IGHG3*\*01/*IGHG1*\*02/*IGHG2*\*06 was common in sub-Saharan African, matching the local ancestries of the each genome(6). The allele population distributions of *IGHG4*, *IGHA1*, and *IGHA2* remained to be investigated.

**Figure 7.**
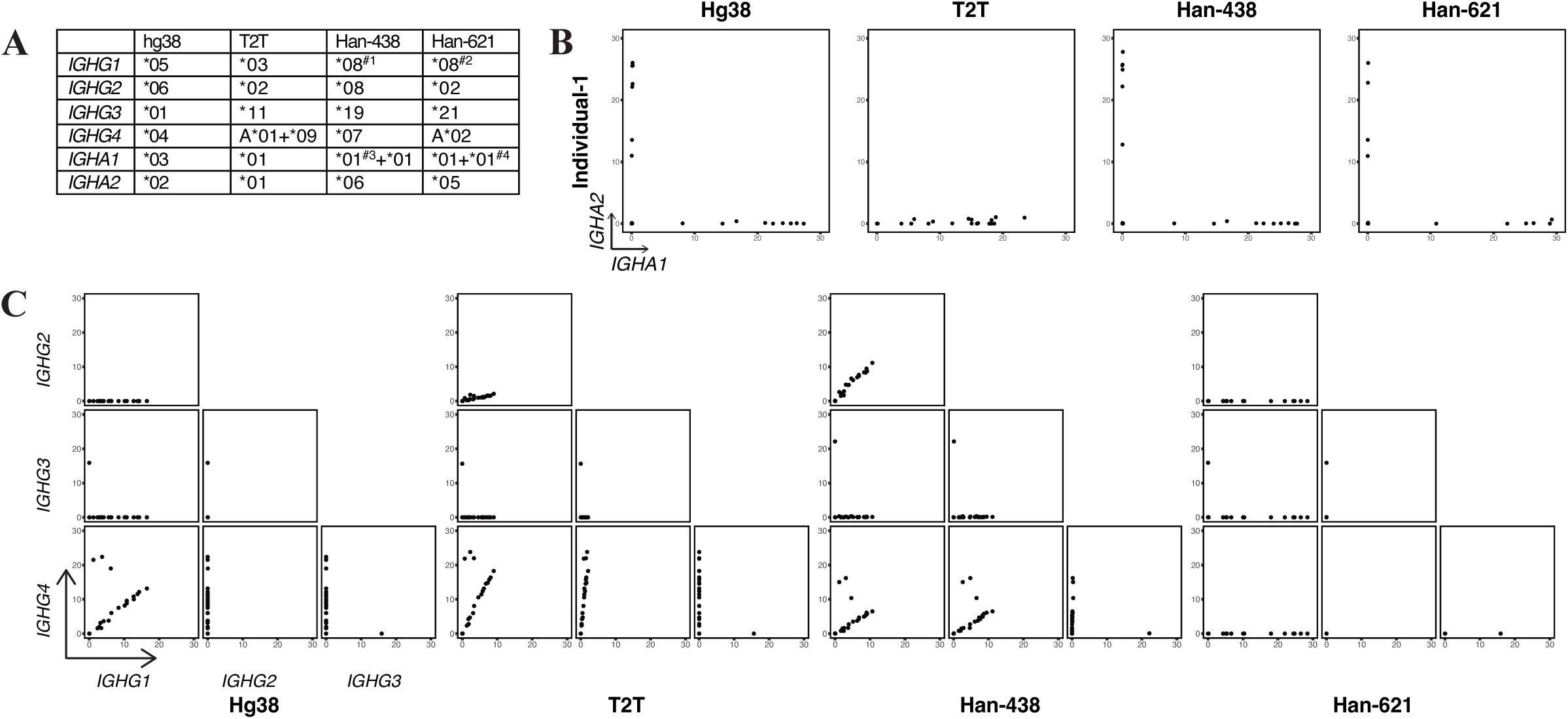
The allele differences between genome references and their impact on isotype calling. **A)** The table shows the IMGT alleles of isotype genes in 4 genome references. #1, there is 1 mismatch in the protein coding region and 3 in the 3-UTR; #2, there are 5 mismatches in the 3-UTR; #3, *IGHA1* is duplicated in Han-438, and for the first *IGHA1*, there is 1 mismatch; and #4, *IGHA1* is duplicated in Han-621, and for the second *IGHA1*, there is 1 mismatch. **B)** Scatter plots correlating the expression levels of *IGHA1* and *IGHA2* for intestinal mucosa ASCs of the first Chinese individual. The genome references used of each plot are shown accordingly. Here expression is represented as reads per hundred (RPH). **C)** Paired scatter plots correlating the expression levels of pairs of IgG subclass genes for intestinal mucosa ASCs of the first Chinese individual. The genome references used of each plot are shown accordingly. Here expression is represented as reads per hundred (RPH).

The genetic background of the in-house BMPC dataset is European Australian. While such information was lacking for the other three non-Chinese public datasets, it is reasonable to assume that they are also of European origin, based on the similar isotype calling outcomes and the fact that the dataset samples were collected in the USA and in Britain. Thus, it was reasonable that the European-origin datasets yielded better results when using T2T as the genome reference. Conversely, mapping the European-origin datasets to the African-origin hg38 gave rise to false results that did not obey the “one cell, one antibody” dogma.

In Bashirova et al.(51), no haplotype analysis was performed for an East Asia population, but they did show that the most frequent alleles of each single gene for East Asia were different from the European or African alleles. To investigate whether T2T also performs better for East Asian data, an additional 10x Genomics Chromium single-cell 3’ dataset including two Han Chinese individuals was examined(37). The 10x data for the two Chinese individuals was mapped to hg38 and T2T, and also to two Han Chinese Han genomes from Liao et al.(40). Han-438, and Han-621, display alternative alleles compared with hg38 and T2T (Fig 7A). With the same quality control and downstream analysis process, the isotype calling was visualized by scatter plots (Fig 7BC, Fig S21).

For IGHA genes, with the “one cell, one antibody” as a ground truth, hg38, Han-438, and Han-621 all displayed good IGHA isotype calling (Fig 7B and S21B). In contrast the T2T output of individual-1 only displayed *IGHA1*^+^ cells, missing *IGHA2* expression and the T2T output of individual-2 displayed *IGHA* double-positive cells.

For IGHG genes, neither hg38, T2T nor Han-438 provided accurate immunoglobulin isotype calling, with all displaying IGHG double-positive cells (Fig 7C and S21C). Only Han-621 provided a relatively better output.

It remains challenging to interrogate the isotypes for East Asians due to the rareness of public datasets including ASCs highly expressing isotype genes but also a lack of a complete and representative East Asian genome assembly. Based on the preliminary analysis of these two Chinese individuals, it can be deduced that T2T is not always the optimal reference genome depending on the ethnicity of the subjects.

## Discussion

While model organisms, such as laboratory mice, have very uniform genetic backgrounds, human individuals have very diverse genetic backgrounds. Despite this known sequence diversity, hg38 was the only human genome reference in standard use until the relatively recent publication of T2T. Whether the variable human genetic backgrounds impact the outcome of sequencing data analysis with a single genome reference remains unknown. Here, our analysis indicated that two genome references, hg38 and T2T with different local ancestries, provided different results in IGH isotype calling for single ASC RNAseq data.

“One cell one antibody” is a cornerstone of the clonal selection theory that has been central to our understanding of the immune system for over 60 years(52). This theory can be employed as a ground truth, where one single cell should only express one isotype gene. When analysing the four public datasets, the T2T output fits this theory, while the widely used hg38 output does not, prompted us to explore the underlying cause of this uncertainty.

The mapping performance of hg38 and T2T were similar on a genome wide level, with the notable exception of the immunoglobulin isotype genes. The hg38 outputs displayed many *IGHG1*^+^*IGHG2*^+^, *IGHG1*^+^*IGHG3*^+^, and *IGHA1*^+^*IGHA2*^+^ isotype double-positive cells. It was unlikely that the dual expression was the result of cell doublets being sequenced together, as these events should be randomly distributed between any two isotypes, instead of just between certain isotypes, as observed. In the T2T outputs, the cells were all single-positive, apart from a negligible number of doublet cells. The cells that were *IGHG1*^+^*IGHG2*^+^ in the hg38 output became *IGHG2*^+^, cells that were *IGHG1*^+^*IGHG3*^+^ became *IGHG1*^+^, and cells that were *IGHA1*^+^*IGHA2*^+^ became *IGHA2*^+^. The conversions were due to the correct assignments of reads that were mapped to certain regions of the 3’ exons (Fig 3C, Fig S11C, and Fig 5C).

In addition, because the sequences of IGHG gene family are more distinguishable in T2T than hg38, many reads that were multimapped in hg38 could be assigned to *IGHG1* and *IGHG2* in the T2T outputs (Fig 4). However, the situation for *IGHA* genes was reversed. Due to a large segment of homologous sequences between *IGHA1* and *IGHA2* in T2T, reads mapped to this segment were regarded as multi-mapping reads and were discarded in the T2T output (Fig 5F). We showed that those discarded reads could be rescued with a modified reference (Fig 6).

Next, we elaborated why the isotype sequences were different between these two genome references. T2T is the first published complete human genome employing the long read sequencing technology(5). T2T is sequenced from a homozygous complete hydatidiform mole (CHM) cell line, representing a complete haplotype from a single European individual(5). In comparison, the current hg38 genome is derived from multiple individuals, with one individual with African-European admixed ancestry dominating(4). The alleles of *IGHG1*, *IGHG2*, and *IGHG3* of T2T and hg38 are distinct and represent the most frequent haplotypes of European and sub-Saharan African ancestry, respectively(51). Thus, it is reasonable that the isotype sequence differences are reflected by the ethnic discrepancy. The ethnicity of the BMPC data is known to us and is of European origin. For the other three non-Chinese public datasets, ethnicity information was not released, but it seems likely that they are also European origin because they have similar isotype calling results to the BMPC data. It could be speculated that mapping European samples to T2T results in better outcomes than mapping to hg38, which reflects a different ancestral genetic background. To address the impact of the genetic background of the genome reference, we introduced one more public dataset from China and demonstrated that neither the European-based T2T nor the African-based hg38 performed better for Chinese samples than the native Han genome references.

Considering the importance of immunoglobulin isotypes in immune regulation and diseases, and also in the monoclonal antibody production industry, it is necessary to identify the isotypes correctly. Based on this study, when using the mapping-count method with a single genome reference, T2T genome reference is a better option than hg38 for European-origin sample but not for East Asian samples. In each circumstance, the genetic background of genome reference will impact the results of sequencing data analysis, at least for the isotype calling, with other gene families remaining to be explored, especially for those that are highly variable between human populations.

The aim of the T2T genome project was to fill in gaps that were present in previous drafts of the human genome, particularly in complex regions such as centromeres and telomeres. The T2T also provides a genome reference with European genetic background. Our study shows that the improvements offered by the T2T genome are not limited to the newly completed regions but, for European samples, are also present in terms of improved accuracy for a protein-coding gene family that was fully covered in older genomes. Our study is limited to immunoglobin genes because it is only for that family that the clonal selection hypothesis of one-cell-one-antibody allows us to identify and diagnose read alignment inaccuracies. There is no reason to expect that sequencing bias should be limited to this one gene family, but the isotype results may be especially sensitive to such biases because of the high level of sequence similarity between the isotype subclasses genes.

## Data Availability

The datasets analysed during the current study are available from the NCBI Sequence Read Archive experiments https://www.ncbi.nlm.nih.gov/sra/SRX20001597, https://www.ncbi.nlm.nih.gov/sra/SRX6584269, https://www.ncbi.nlm.nih.gov/sra/SRX9986858 and https://www.ncbi.nlm.nih.gov/sra/SRX4720060.

Computer code used to generate results in this manuscript is provided in Supplementary Materials.

## Supporting information

Supplementary figures, table and computer code

## Acknowledgements

Matthew Robertson from 10X genomics provided technical support with regard to custom reference making and BAM file dissection.

