## Supplementary figures, table and computer code for "T2T-CHM13 vs GRCh38: accurate identification of immunoglobulin isotypes from scRNA-seq requires a genome reference matched for ethnicity"

### Figure S1

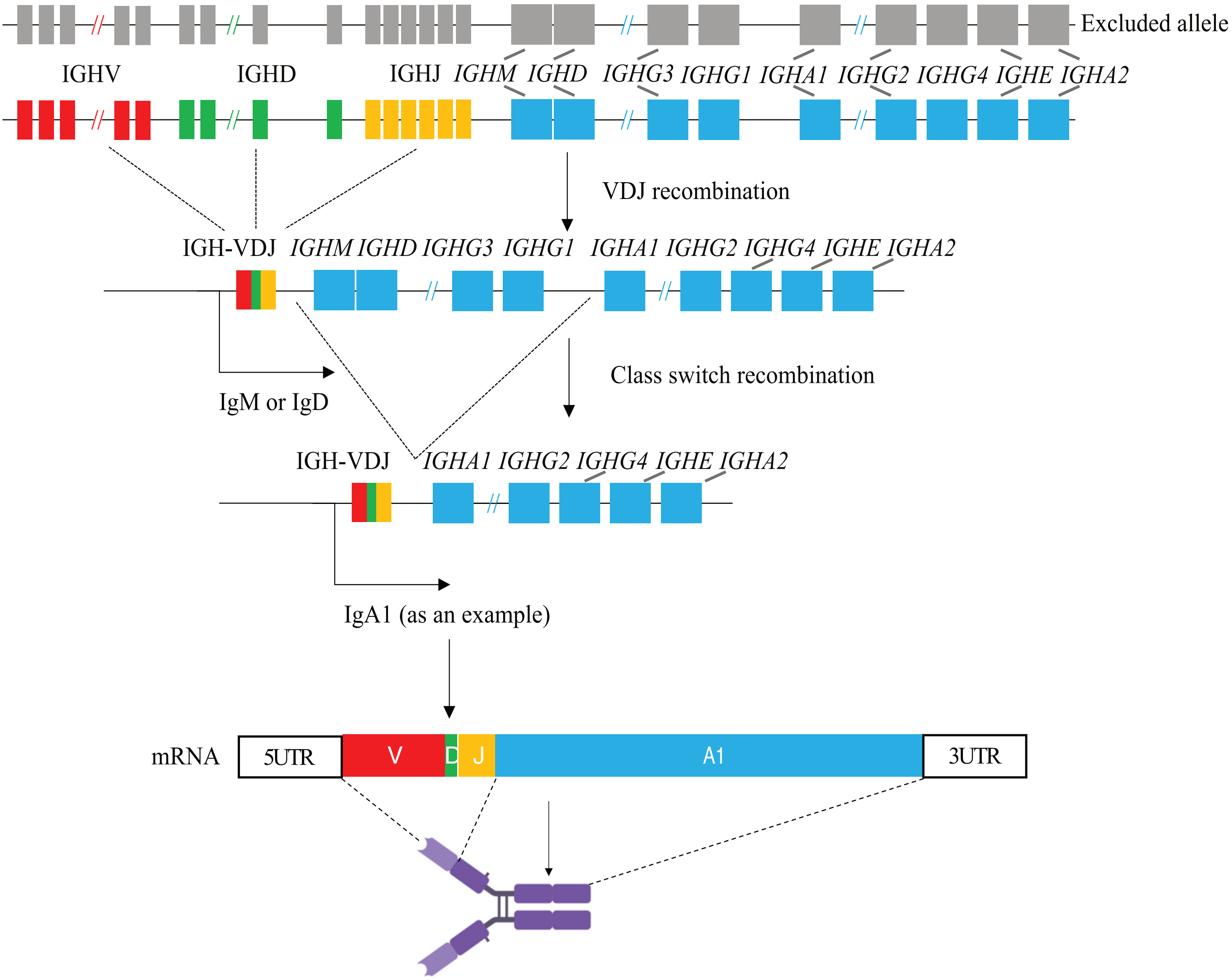


**Figure S1 Somatic recombination of the *IGH* locus in human B cells**

Schematic plots showing the structure of the human immunoglobulin heavy chain gene (*IGH*) and the process of VDJ recombination and class switch recombination. The first row shows the structure of *IGH* gene locus in germ line state, where red rectangles represent *IGHV* genes, green rectangles represent *IGHD* genes, and blue rectangles represent *IGHC* genes. The second row shows the structure of *IGH* gene locus after VDJ recombination during B cell development in which a single VDJ exon is synthesised using one *IGHV* gene, one *IGHD* gene and one *IGHJ* gene while the IGHC genes remain unchanged. At this stage, the VDJ exon will splice to *IGHM* or *IGHD* by alternative splicing to generate IgM or IgD. Once B cells become activated, class switch recombination is performed, after which the DNA sequence between VDJ exon and any *IGHC* gene except *IGHM* and *IGHD* (*IGHA1* is shown as an example) will be removed as shown in the bottom row. After class switch recombination, the VDJ exon will splice to the most 5' remaining IGHC, in this case *IGHA1* to generate IgA1. The structure of mRNA and its protein product heavy chains (in the dimer state) are shown in the bottom. "//" in red or green means that some *IGHV* genes or *IGHD* genes are not shown, while "//" in blue means that a long DNA segment is omitted due to space limitations.

### Figure S2

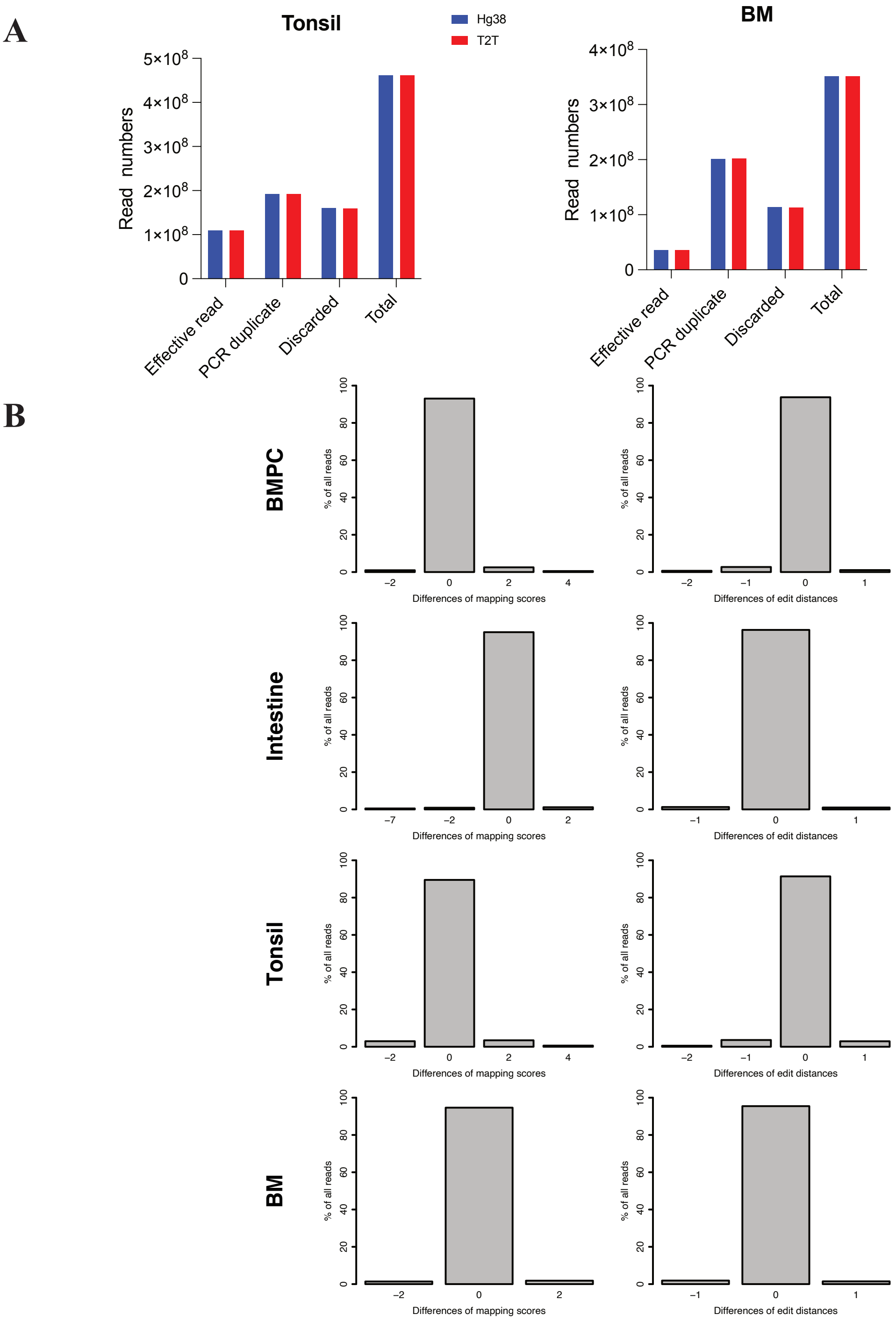


**Figure S2 Mapping performance is comparable between hg38 and T2T**

**A)** Classification of reads for the tonsil (left) and bone marrow (right) datasets for each reference genome. The barplots show the total read count for each sample and also the number of effective reads, the number of reads identified as PCR duplicates and the number of reads discarded for other reasons. Only effective reads were used for downstream analyses.

**B)** Differences in mapping scores (left, extracted from AS tag of the BAM file) and edit distances (right, extracted from nM tag of the BAM file). For each sequence read, the T2T mapping score or edit distance was subtracted from the hg38 mapping score or edit distance for the same read. The left panels show the percentage of reads with each mapping score difference. The right panels are similar but for differences in edit distances. Only values with >0.5% are shown. Rows correspond to the four different datasets. Most reads show 0 difference, meaning that the mapping performance for most of the reads is comparable.

### Figure S3

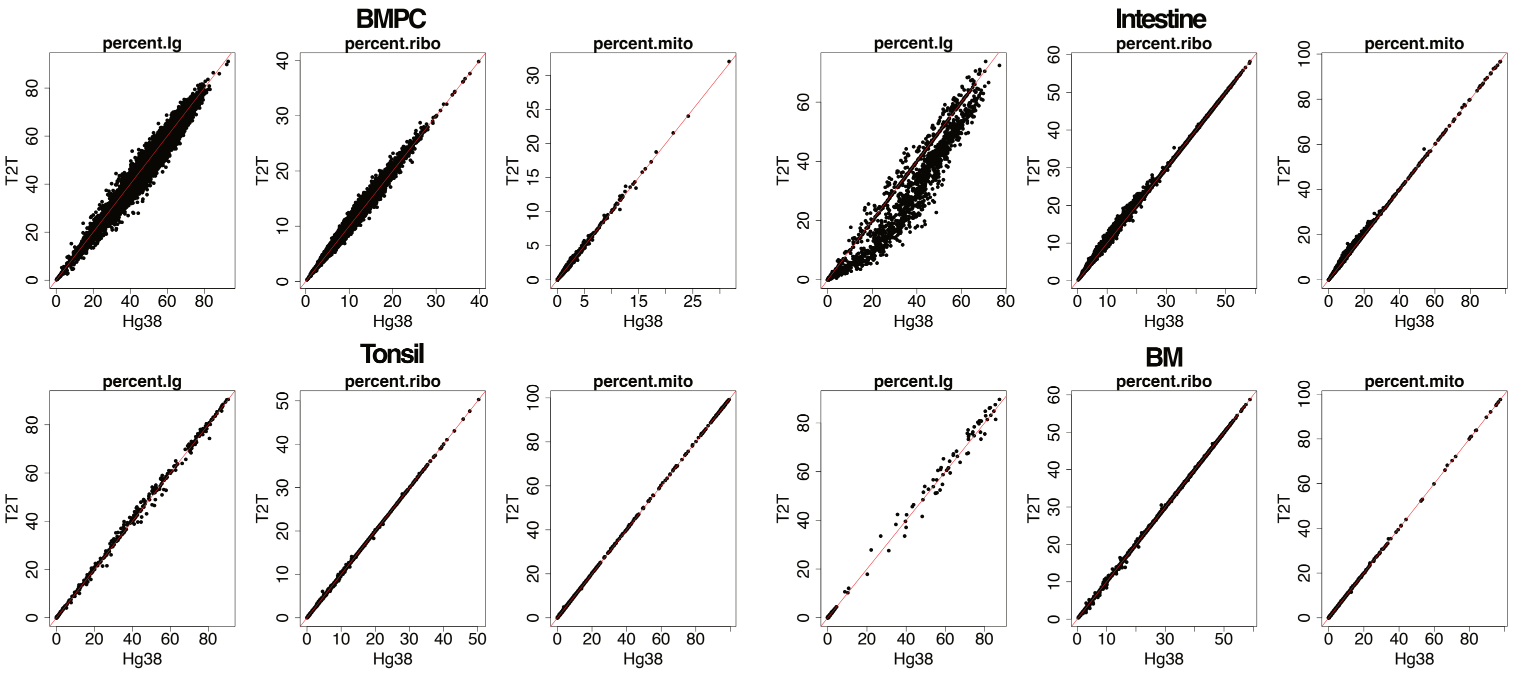


**Figure S3 The quality control**

The metrics (the left of each panel, the percentages of expressed immunoglobulin genes; the middle of each panel, the percentages of expressed ribosomal genes; and the right of each panel, the percentages of expressed mitochondrial genes) of the 4 datasets used for quality control. The hg38 and T2T cell-wise metrics are compared before quality control by scatter plots. Red lines in scatter plots represent the x=y line.

### Figure S4

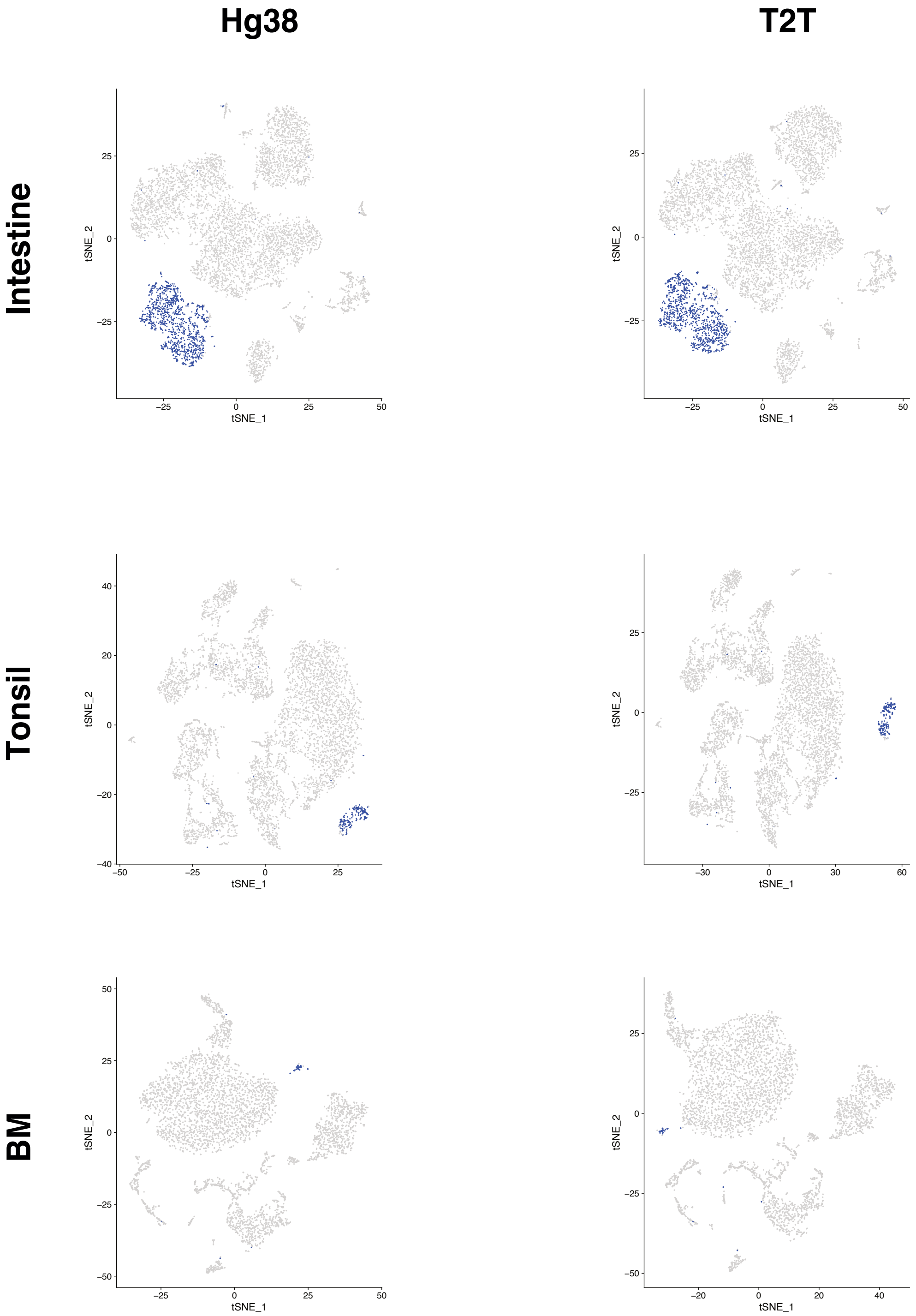


**Figure S4 Cell clustering is comparable between hg38 and T2T**

t-SNE plots of all cells for the intestinal mucosa (top), tonsil (middle) and bone marrow (bottom) datasets for each reference genome. Antibody-secreting cells (ASCs, highlighted in blue) cluster separately from other cells and were selected for downstream isotype analyses. (The BMPC dataset, was already sorted for ASCs before scRNA-seq profiling, so all cells for that dataset were shown in Figure 2A.)

### Figure S5

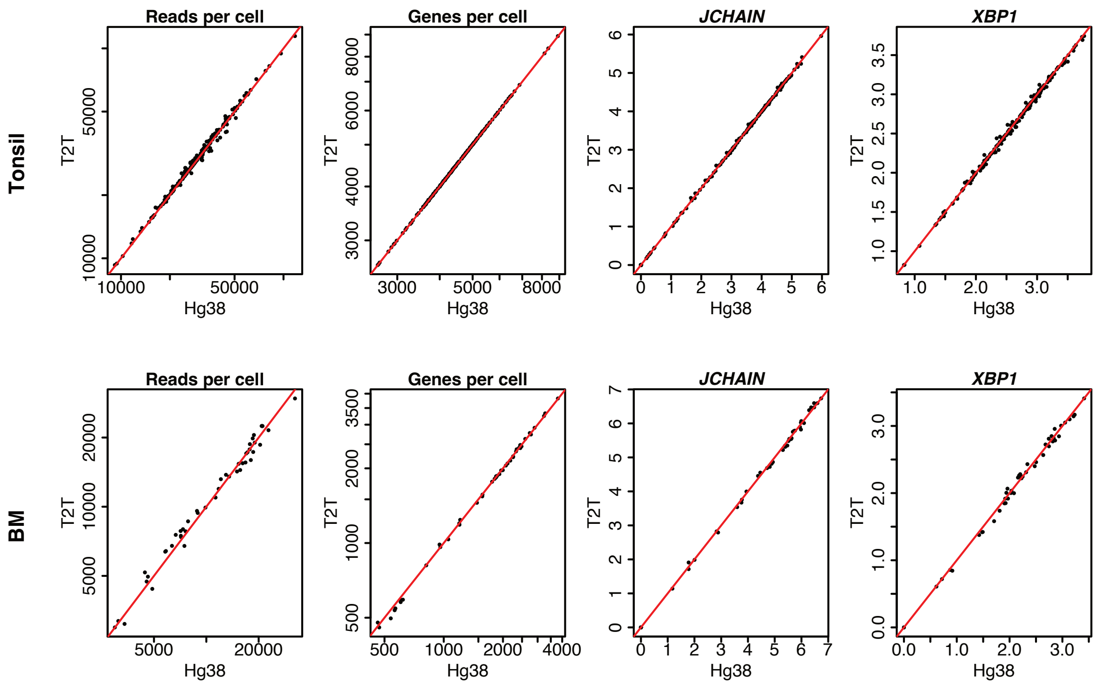


**Figure S5 Cellular metrics for quality control and key gene expression are comparable between hg38 and T2T**

Reads per cell, genes detected per cell, *JCHAIN* expression, and *XBP1* expression for the tonsil (top) and bone marrow (bottom) datasets. Here and subsequently, only bona fide ASCs are shown. The hg38 and T2T cell-wise counts and expression value are compared by scatter plots. Red lines in scatter plots represent the x=y line.

### Figure S6

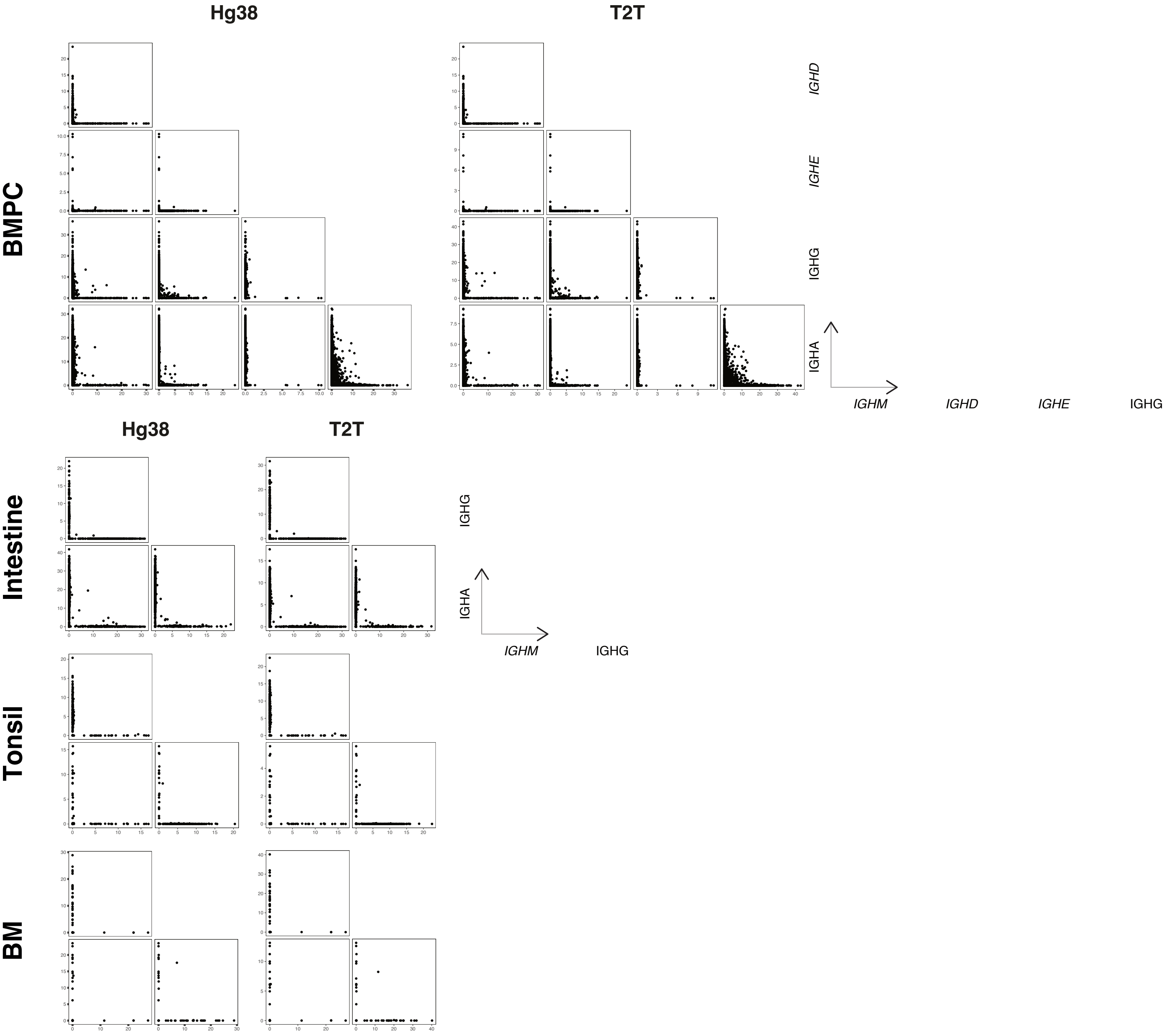


**Figure S6 The isotype gene expression**

Paired scatter plots correlating the expression levels of pairs of *IGHM*, *IGHD*, *IGHE*, total IGHG, and total IGHA for the 4 datasets. Plots are shown for hg38 (left) and T2T (right). Expression is represented as reads per hundred (RPH). In some datasets, the expression of *IGHD* and *IGHE* is omitted.

### Figure S7

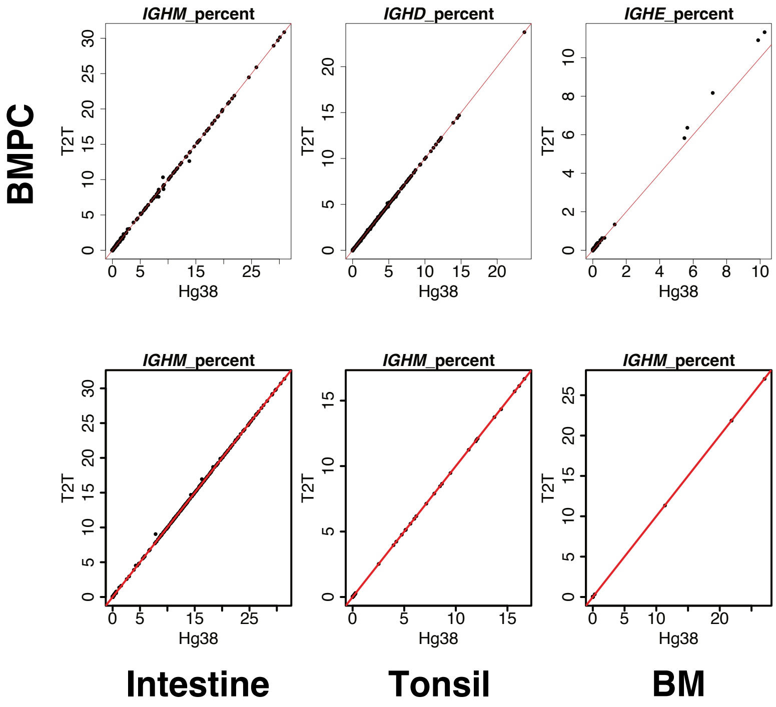


**Figure S7 The comparison of *IGHM*, *IGHD*, and *IGHE* expression between hg38 and T2T**

Top, *IGHM* (left), *IGHD* (middle), and *IGHE* (right) expression for the BMPC datasets. Bottom, *IGHM* expression for the remaining 3 datasets. The hg38 and T2T cell-wise counts are compared by scatter plots. Red lines in scatter plots represent the x=y line.

### Figure S8

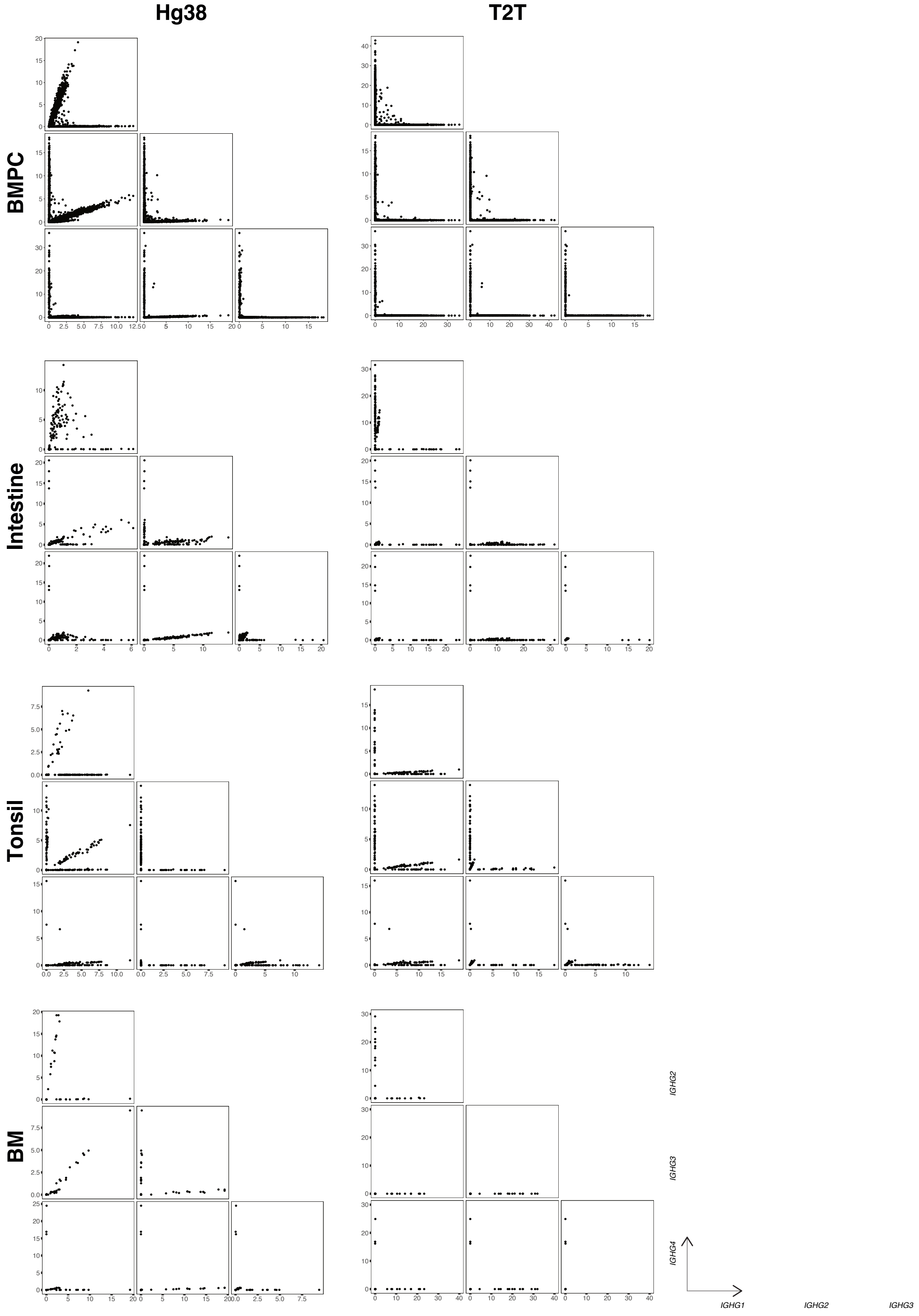


**Figure S8 The T2T genome identifies single IgG subclasses for single cells**

Paired scatter plots correlating the expression levels of pairs of IgG subclass genes for the 4 datasets. Plots are shown for hg38 (left) and T2T (right). Expression is represented as reads per hundred (RPH).

### Figure S9

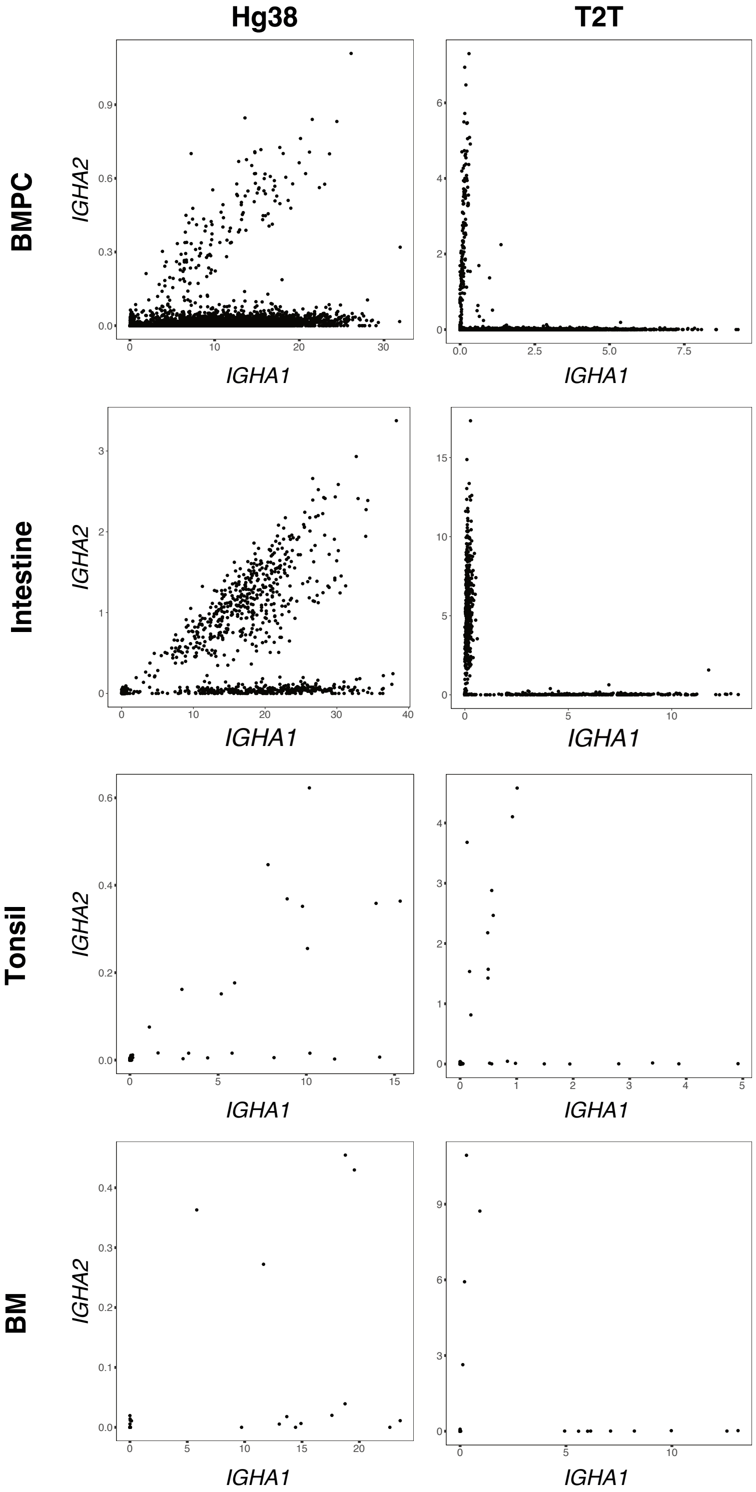


**Figure S9 The T2T genome identifies single IgA subclasses for single cells**

Scatter plots correlating *IGHA1* and *IGHA2* expression for the 4 datasets. Left plots are for hg38 and right plots for T2T. Expression is represented as reads per hundred (RPH).

**Figure S10**

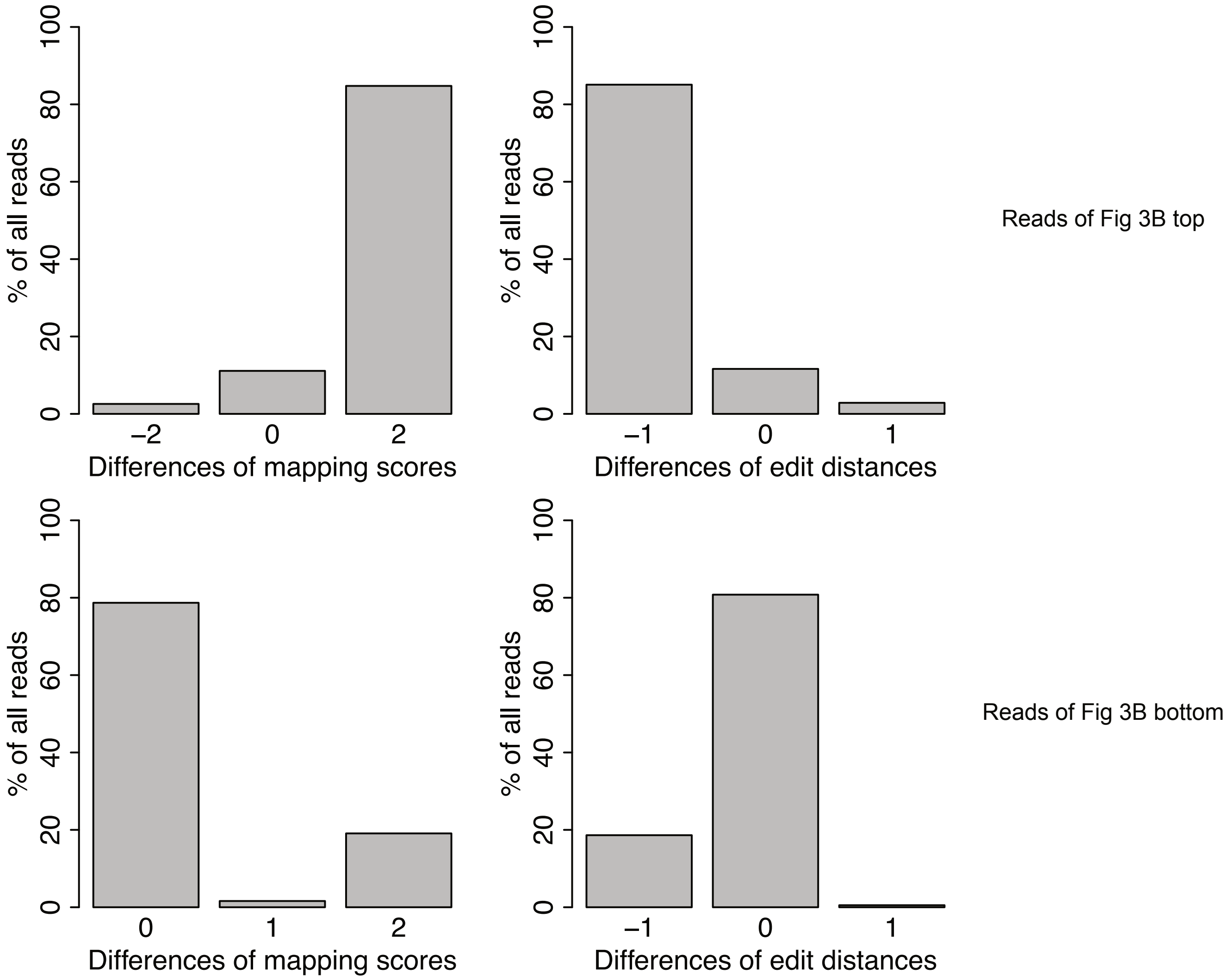


**Figure S10 The mapping performance of IGHG double-positive cells**

Differences in mapping scores (left, extracted from AS tag of the BAM file) and edit distances (right, extracted from nM tags of the BAM file). For each sequence read, the T2T mapping score or edit distance was subtracted from the hg38 mapping score or edit distance for the same read. The left panels show the percentage of reads with each mapping score difference. The right panels are similar but for differences in edit distances. Only values with >0.5% are shown. The top row corresponds to the reads of Fig 3B top and the bottom row corresponds to the reads of Fig 3B bottom. 0 difference means that the mapping performance is comparable between T2T and hg38; positive difference for mapping scores or negative difference for edit distances means that the mapping performance of T2T is better than hg38; and *vice versa*.

### Figure S11

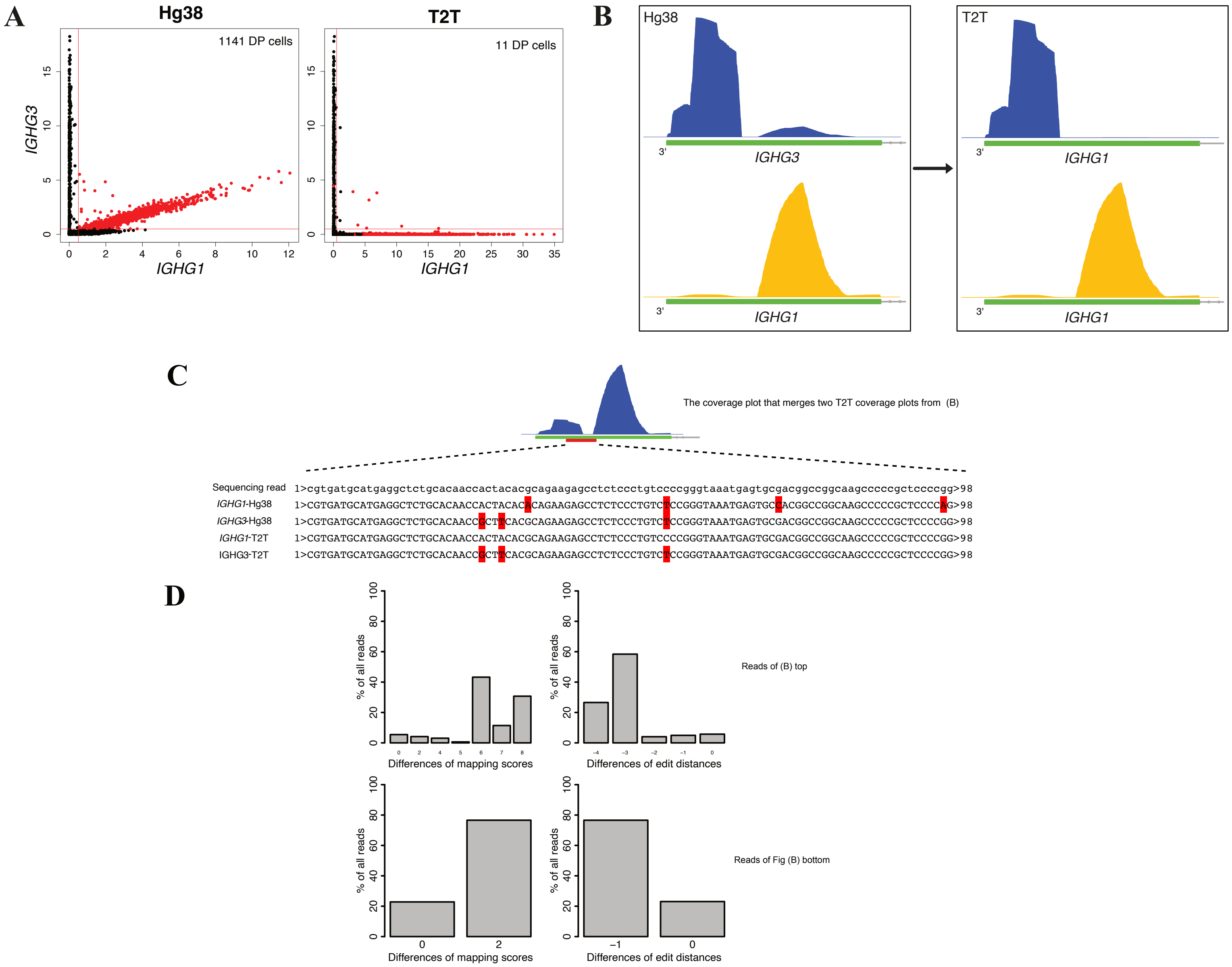


**Figure S11 Separating *IGHG1*⁺ and *IGHG3*⁺ plasma cells**

**A)** Scatter plots correlating *IGHG1* and *IGHG3* expression for BMPC. Left plot for hg38 and right plot for T2T. The *IGHG1*+*IGHG3*+ cells of the hg38 output are highlighted in both plots. The horizontal and vertical red lines represent the thresholds for positive cells of an isotype gene. Here, the thresholds are 0.5% for both *IGHG1* and *IGHG3*. The numbers of double-positive (DP) cells for both outputs are also shown.

**B)** Coverage plots for reads from the highlighted cells in (A). Top left plot shows reads mapped to *IGHG3* using hg38 and bottom left shows reads mapped to *IGHG1* using hg38. Only the CH3-CHS exon of the genes are shown. The plots on the right show coverage plots for the same reads but using T2T. The top right plot shows that all the reads mapped to *IGHG3* using hg38 map instead to *IGHG1* using T2T. The bottom right plot shows that all reads mapped to *IGHG1* using hg38 map also map to *IGHG1* using T2T.

**C)** Alignment results for one illustrative sequence read incorrectly assigned to *IGHG3* by hg38 in (B). The full sequence of the read is shown together with the best matches to the *IGHG1*-hg38, *IGHG3*-hg38, *IGHG1*-T2T and *IGHG3*-T2T genome regions. Mismatched bases are marked in red. The read maps perfectly to *IGHG1*-T2T and, using T2T, the assignment to *IGHG1* is unambiguous. Using hg38, however, the result is reversed with fewer mismatches for *IGHG3*-hg38 than for *IGHG1*-hg38. The coverage plot merges the right two plots from (B) on the same scale, i.e., shows *IGHG1*-T2T coverage for all reads mapped by hg38 to either *IGHG1* or *IGHG3*. The aligned position of the read is shown by a red bar.

**D)** Differences in mapping scores (left, extracted from AS tag of the BAM file) and edit distances (right, extracted from nM tags of the BAM file). For each sequence read, the T2T mapping score or edit distance was subtracted from the hg38 mapping score or edit distance for the same read. The left panels show the percentage of reads with each mapping score difference. The right panels are similar but for differences in edit distances. Only values with >0.5% are shown. The top row corresponds to the reads of (B) top and the bottom row corresponds to the reads of (B) bottom. 0 difference means that the mapping performance is comparable between T2T and hg38; positive difference for mapping scores or negative difference for edit distances means that the mapping performance of T2T is better than hg38; and *vice versa*.

### Figure S12

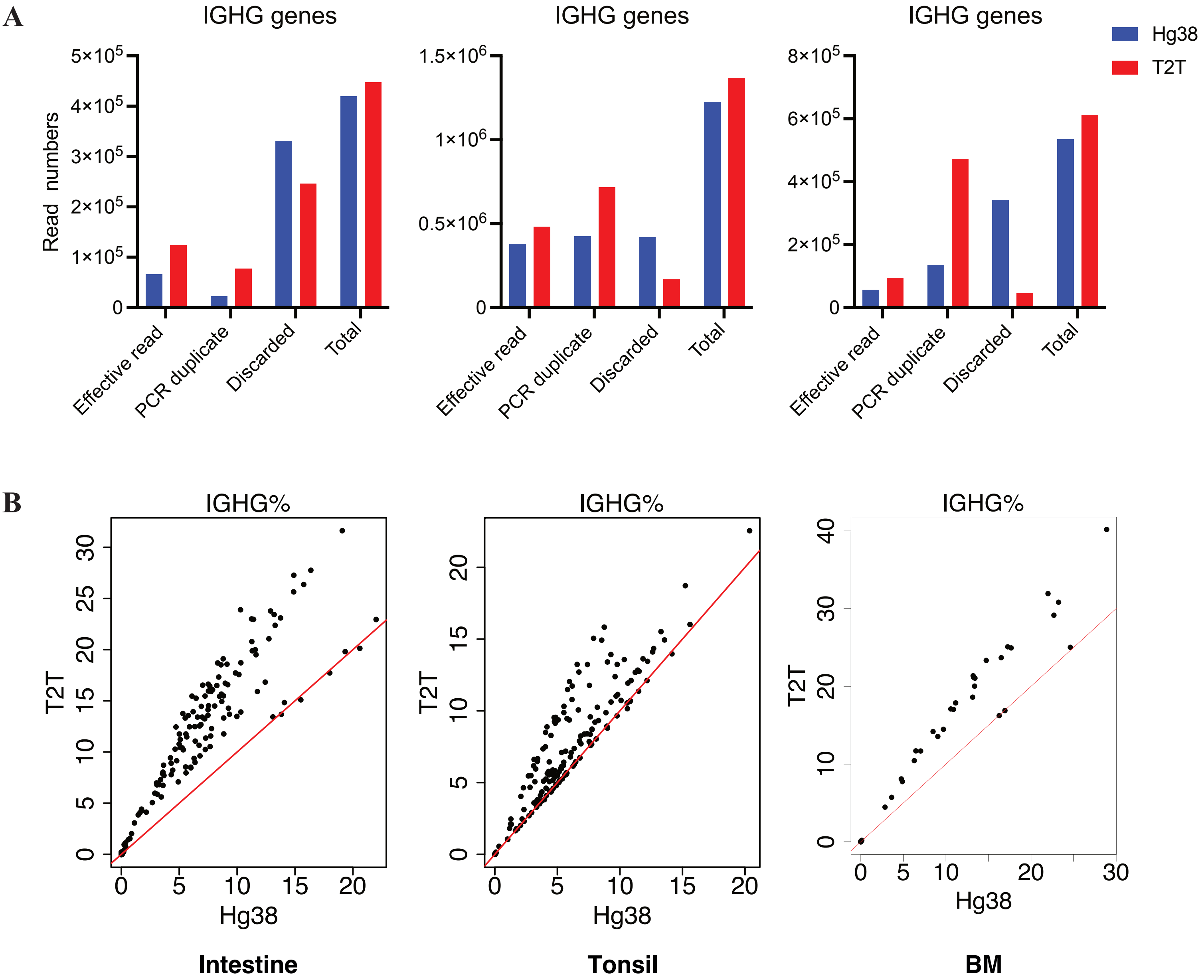


**Figure S12 T2T has more reads effectively mapped to IGHG genes than hg38**

**A)** Classification of reads that are assigned to IGHG genes for the intestine (left), tonsil (middle) and bone marrow (right) datasets for each reference genome. The barplots show the total read count of IGHG genes and also the number of effective reads, the number of reads identified as PCR duplicates and the number of reads discarded for other reasons.

**B)** Percentage of IGHG gene reads per cell for the intestine (left), tonsil (middle) and bone marrow (right) datasets. Hg38 and T2T results are compared by scatter plots. Red lines in scatter plots represent the x=y line.

### Figure S13

**A**

```
                          *         *         *         *         *         *         *         *         *         *
IGHG1-T2T      1>GGCAGCCCCGAGAACCACAGGTGTACACCCTGCCCCCATCCCGGGAGGAGATGACCAAGAACCAGGTCAGCCTGACCTGCCTGGTCAAAGGCTTCTATCC>100
IGHG2-T2T      1>GGCAGCCCCGAGAACCACAGGTGTACACCCTGCCCCCATCCCGGGAGGAGATGACCAAGAACCAGGTCAGCCTGACCTGCCTGGTCAAAGGCTTCTACCC>100
IGHG3-T2T      1>GACAGCCCCGAGAACCACAGGTGTACACCCTGCCCCCATCCCGGGAGGAGATGACCAAGAACCAGGTCAGCCTGACCTGCCTGGTCAAAGGCTTCTACCC>100
IGHG4-1-T2T    1>GGCAGCCCCGAGAGCCACAGGTGTACACCCTGCCCCCATCCCAGGAGGAGATGACCAAGAACCAGGTCAGCCTGACCTGCCTGGTCAAAGGCTTCTACCC>100
IGHG4-2-T2T    1>GGCAGCCCCGAGAGCCACAGGTGTACACCCTGCCCCCATCCCAGGAGGAGATGACCAAGAACCAGGTCAGCCTGACCTGCCTGGTCAAAGGCTTCTACCC>100

                          *         *         *         *         *         *         *         *         *         *
             101>CAGCGACATCGCCGTGGAGTGGGAGAGCAATGGGCAGCCGGAGAACAACTACAAGACCACGCCTCCCGTGCTGGACTCCGACGGCTCCTTCTTCCTCTAT>200
             101>CAGCGACATCGCCGTGGAGTGGGAGAGCAATGGGCAGCCGGAGAACAACTACAAGACCACACCTCCCATGCTGGACTCCGACGGCTCCTTCTTCCTCTAC>200
             101>CAGCGACATCGCCGTGGAGTGGGAGAGCAGCGGGCAGCCGGAGAACAACTACAACACCACGCCTCCCATGCTGGACTCCGACGGCTCCTTCTTCCTCTAC>200
             101>CAGCGACATCGCCGTGGAGTGGGAGAGCAATGGGCAGCCGGAGAACAACTACAAGACCACGCCTCCCGTGCTGGACTCCGACGGCTCCTTCTTCCTCTAC>200
             101>CAGCGACATCGCCGTGGAGTGGGAGAGCAATGGGCAGCCGGAGAACAACTACAAGACCACGCCTCCCGTGCTGGACTCCGACGGCTCCTTCTTCCTCTAC>200

                          *         *         *         *         *         *         *         *         *         *
             201>AGCAAGCTCACCGTGGACAAGAGCAGGTGGCAGCAGGGGAACGTCTTCTCATGCTCCGTGATGCATGAGGCTCTGCACAACCACTACACGCAGAAGAGCC>300
             201>AGCAAGCTCACCGTGGACAAGAGCAGGTGGCAGCAGGGGAACGTCTTCTCATGCTCCGTGATGCATGAGGCTCTGCACAACCACTACACACAGAAGAGCC>300
             201>AGCAAGCTCACCGTGGACAAGAGCAGGTGGCAGCAGGGGAACATCTTCTCATGCTCCGTGATGCATGAGGCTCTGCACAACCGCTTCACGCAGAAGAGCC>300
             201>AGCAGGCTAACCGTGGACAAGAGCAGGTGGCAGGAGGGGAATGTCTTCTCATGCTCCGTGATGCATGAGGCTCTGCACAACCACTACACGCAGAAGAGCC>300
             201>AGCAAGCTCACCGTGGACAAGAGCAGGTGGCAGGAGGGGAATGTCTTCTCATGCTCCGTGATGCATGAGGCTCTGCACAACCACTACACACAGAAGAGCC>300

                          *         *         *         *         *         *         *         *         *         *
             301>TCTCCCTGTCCCCGGGTAAATGAgtgcgacggccggcaagcccccgctccccgggctctcgcggtcgcacgaggatgcttggcacgtaccccgtctacat>400
             301>TCTCCCTGTCTCCGGGTAAATGAgtgccacggccagcaagcccccgctccccaggctctcgggtcgcgcgaggatgcttggcacgtaccccgtgtacat>400
             301>TCTCCCTGTCTCCGGGTAAATGAgtgcgacggccggcaagcccccgctccccgggctctcgggtcgcgcgaggatgcttggcacgtaccccgtgtacat>400
             301>TCTCCCTGTCTCTGGGTAAATGAgtgccaggccggcaagcccccgctccccgggctctcgggtcgcgcgaggatgcttggcacgtaccccgtgtacat>400
             301>TCTCCCTGTCTCTGGGTAAATGAgtgccaggccggcaagcccccgctccccgggctctcgggtcgcgcgaggatgcttggcacgtaccccgtctacat>400

                          *         *         *         *         *
             401>acttcccaggcacccagcatggaaataaagcacccaccactgccctgggcccctgcga>458
             401>acttcccgggcacccagcatggaaataaagcacccagcgcttccctgggcccctgcga>458
             401>acttcccgggcacccagcatggaaataaagcacccagcgctgccctgggcccctgcaa>458
             401>acttcccgggcgcccagcatggaaataaagcacccagcgctgccctgggcccctgcga>458
             401>acttcccaggcacccagcatggaaataaagcacccaccactgccctgggcccctgtga>458
```

**B**

```
                          *         *         *         *         *         *         *         *         *         *
IGHG1-hg38     1>GGCAGCCCCGAGAACCACAGGTGTACACCCTGCCCCCATCCCGGGATGAGCTGACCAAGAACCAGGTCAGCCTGACCTGCCTGGTCAAAGGCTTCTATCC>100
IGHG2-hg38     1>GGCAGCCCCGAGAACCACAGGTGTACACCCTGCCCCCATCCCGGGAGGAGATGACCAAGAACCAGGTCAGCCTGACCTGCCTGGTCAAAGGCTTCTACCC>100
IGHG3-hg38     1>GACAGCCCCGAGAACCACAGGTGTACACCCTGCCCCCATCCCGGGAGGAGATGACCAAGAACCAGGTCAGCCTGACCTGCCTGGTCAAAGGCTTCTACCC>100
IGHG4-hg38     1>GGCAGCCCCGAGAGCCACAGGTGTACACCCTGCCCCCATCCCAGGAGGAGATGACCAAGAACCAGGTCAGCCTGACCTGCCTGGTCAAAGGCTTCTACCC>100

                          *         *         *         *         *         *         *         *         *         *
             101>CAGCGACATCGCCGTGGAGTGGGAGAGCAATGGGCAGCCGGAGAACAACTACAAGACCACGCCTCCCGTGCTGGACTCCGACGGCTCCTTCTTCCTCTAC>200
             101>CAGCGACATCTCCGTGGAGTGGGAGAGCAATGGGCAGCCGGAGAACAACTACAAGACCACACCTCCCATGCTGGACTCCGACGGCTCCTTCTTCCTCTAC>200
             101>CAGCGACATCGCCGTGGAGTGGGAGAGCAGCGGGCAGCCGGAGAACAACTACAACACCACGCCTCCCATGCTGGACTCCGACGGCTCCTTCTTCCTCTAC>200
             101>CAGCGACATCGCCGTGGAGTGGGAGAGCAATGGGCAGCCGGAGAACAACTACAAGACCACGCCTCCCGTGCTGGACTCCGACGGCTCCTTCTTCCTCTAC>200

                          *         *         *         *         *         *         *         *         *         *
             201>AGCAAGCTCACCGTGGACAAGAGCAGGTGGCAGCAGGGGAACGTCTTCTCATGCTCCGTGATGCATGAGGCTCTGCACAACCACTACACACAGAAGAGCC>300
             201>AGCAAGCTCACCGTGGACAAGAGCAGGTGGCAGCAGGGGAACGTCTTCTCATGCTCCGTGATGCATGAGGCTCTGCACAACCACTACACACAGAAGAGCC>300
             201>AGCAAGCTCACCGTGGACAAGAGCAGGTGGCAGCAGGGGAACATCTTCTCATGCTCCGTGATGCATGAGGCTCTGCACAACCGCTTCACGCAGAAGAGCC>300
             201>AGCAGGCTCACCGTGGACAAGAGCAGGTGGCAGGAGGGGAATGTCTTCTCATGCTCCGTGATGCATGAGGCTCTGCACAACCACTACACACAGAAGAGCC>300

                          *         *         *         *         *         *         *         *         *         *
             301>TCTCCCTGTCTCCGGGTAAATGAgtgccacggccggcaagcccccgctccccaggctctcggggtcgcgcgaggatgcttggcacgtaccccgtgtacat>400
             301>TCTCCCTGTCTCCGGGTAAATGAgtgccacggccggcaagcccccgctccccaggctctcggggtcgcgcgaggatgcttggcacgtaccccgtctacat>400
             301>TCTCCCTGTCTCCGGGTAAATGAgtgcgacggccggcaagcccccgctccccgggctctcggggtcgcgcgaggatgcttggcacgtaccccgtgtacat>400
             301>TCTCCCTGTCTCTGGGTAAATGAgtgccagggccggcaagcccccgctccccgggctctcggggtcgcgcgaggatgcttggcacgtaccccgtgtacat>400

                          *         *         *         *         *
             401>acttcccaggcacccagcatggaaataaagcacccagcgcttccctgggcccctgcga>458
             401>acttcccgggcacccagcatggaaataaagcacccagcgctgccctgggcccctgcga>458
             401>acttcccgggcacccagcatggaaataaagcacccagcgctgccctgggcccctgcga>458
             401>acttcccgggcgcccagcatggaaataaagcacccagcgctgccctgggcccctgcga>458
```

**Figure S13 The sequence comparison of IGHG genes**

**A)** Detailed alignment results for the CH3-CHS exon of all IGHG genes for T2T. Mismatched bases are marked in red. 3-UTR regions are represented in lowercase.

**B)** Detailed alignment results for the CH3-CHS exon of all IGHG genes for hg38. Mismatched bases are marked in red. 3-UTR regions are represented in lowercase.

### Figure S14

**A**

```
                         *         *         *         *         *         *         *         *         *         *
IGHG3-T2T     1>GACAGCCCCGAGAACCACAGGTGTACACCCTGCCCCCATCCCGGGAGGAGATGACCAAGAACCAGGTCAGCCTGACCTGCCTGGTCAAAGGCTTCTACCC>100
IGHG3-Hg38    1>GACAGCCCCGAGAACCACAGGTGTACACCCTGCCCCCATCCCGGGAGGAGATGACCAAGAACCAGGTCAGCCTGACCTGCCTGGTCAAAGGCTTCTACCC>100

                         *         *         *         *         *         *         *         *         *         *
            101>CAGCGACATCGCCGTGGAGTGGGAGAGCAGCGGGCAGCCGGAGAACAACTACAACACCACGCCTCCCATGCTGGACTCCGACGGCTCCTTCTTCCTCTAC>200
            101>CAGCGACATCGCCGTGGAGTGGGAGAGCAGCGGGCAGCCGGAGAACAACTACAACACCACGCCTCCCATGCTGGACTCCGACGGCTCCTTCTTCCTCTAC>200

                         *         *         *         *         *         *         *         *         *         *
            201>AGCAAGCTCACCGTGGACAAGAGCAGGTGGCAGCAGGGGAACATCTTCTCATGCTCCGTGATGCATGAGGCTCTGCACAACCGCTTCACGCAGAAGAGCC>300
            201>AGCAAGCTCACCGTGGACAAGAGCAGGTGGCAGCAGGGGAACATCTTCTCATGCTCCGTGATGCATGAGGCTCTGCACAACCGCTTCACGCAGAAGAGCC>300

                         *         *         *         *         *         *         *         *         *         *
            301>TCTCCCTGTCTCCGGGTAAATGAgtgcgacggccggcaagcccccgctccccgggctctcggggtcgcgcgaggatgcttggcacgtaccccgtgtacat>400
            301>TCTCCCTGTCTCCGGGTAAATGAgtgcgacggccggcaagcccccgctccccgggctctcggggtcgcgcgaggatgcttggcacgtaccccgtgtacat>400

                         *         *         *         *         *
            401>acttcccgggcacccagcatggaaataaagcacccagcgctgccctgggcccctgcaa>458
            401>acttcccgggcacccagcatggaaataaagcacccagcgctgccctgggcccctgcga>458
```

**B**

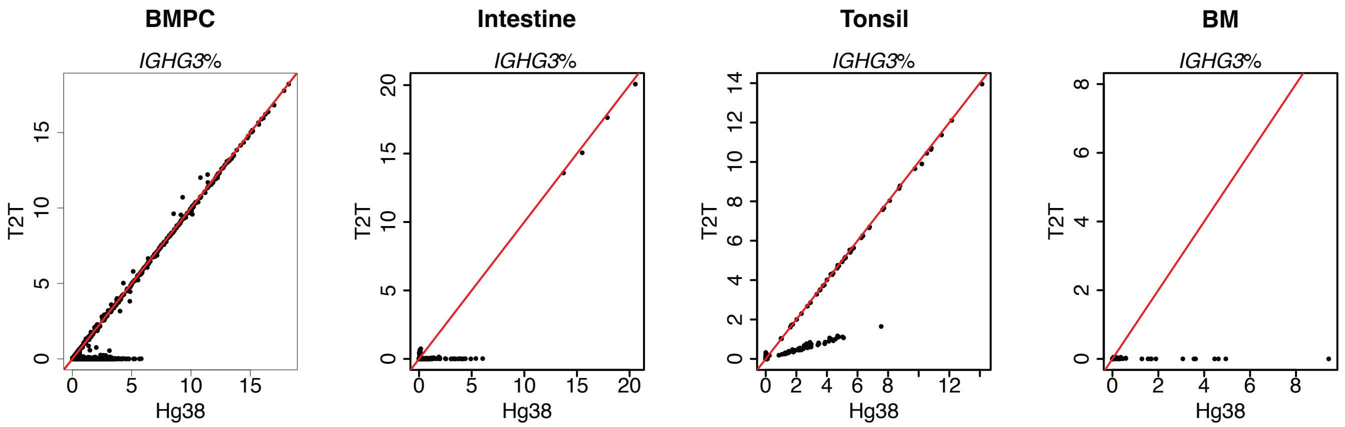


**Figure S14 The comparison of *IGHG3* between T2T and hg38**

**A)** Detailed alignment results for the CH3-CHS exons of *IGHG3* for both T2T and hg38. Mismatched bases are marked in red. 3-UTR regions are represented in lowercase.

**B)** Percentage of *IGHG3* reads per cell for the BMPC, intestine, tonsil and bone marrow datasets. Hg38 and T2T results are compared by scatter plots. Red lines in scatter plots represent the x=y line.

### Figure S15

**A**

```
IGHG1-T2T    1>GGCAGCCCCGAGAACCACAGGTGTACACCCTGCCCCCATCCCGGGAGGAGATGACCAAGAACCAGGTCAGCCTGACCTGCCTGGTCAAAGGCTTCTATCC>100
IGHG1-hg38   1>GGCAGCCCCGAGAACCACAGGTGTACACCCTGCCCCCATCCCGGGATGAGCTGACCAAGAACCAGGTCAGCCTGACCTGCCTGGTCAAAGGCTTCTATCC>100

           101>CAGCGACATCGCCGTGGAGTGGGAGAGCAATGGGCAGCCGGAGAACAACTACAAGACCACGCCTCCCGTGCTGGACTCCGACGGCTCCTTCTTCCTCTAT>200
           101>CAGCGACATCGCCGTGGAGTGGGAGAGCAATGGGCAGCCGGAGAACAACTACAAGACCACGCCTCCCGTGCTGGACTCCGACGGCTCCTTCTTCCTCTAC>200

           201>AGCAAGCTCACCGTGGACAAGAGCAGGTGGCAGCAGGGGAACGTCTTCTCATGCTCCGTGATGCATGAGGCTCTGCACAACCACTACACGCAGAAGAGCC>300
           201>AGCAAGCTCACCGTGGACAAGAGCAGGTGGCAGCAGGGGAACGTCTTCTCATGCTCCGTGATGCATGAGGCTCTGCACAACCACTACACACAGAAGAGCC>300

           301>TCTCCCTGTCCCCGGGTAAATGAgtgcgacggccggcaagcccccgctccccgggctctcgcggtcgcacgaggatgcttggcacgtaccccgtctacat>400
           301>TCTCCCTGTCTCCGGGTAAATGAgtgccacggccggcaagcccccgctccccaggctctcgggtcgcgcgaggatgcttggcacgtaccccgtgtacat>400

           401>acttcccaggcacccagcatggaaataaagcacccaccactgccctgggcccctgcga>458
           401>acttcccaggcacccagcatggaaataaagcacccagcgcttccctgggcccctgcga>458
```

**B**

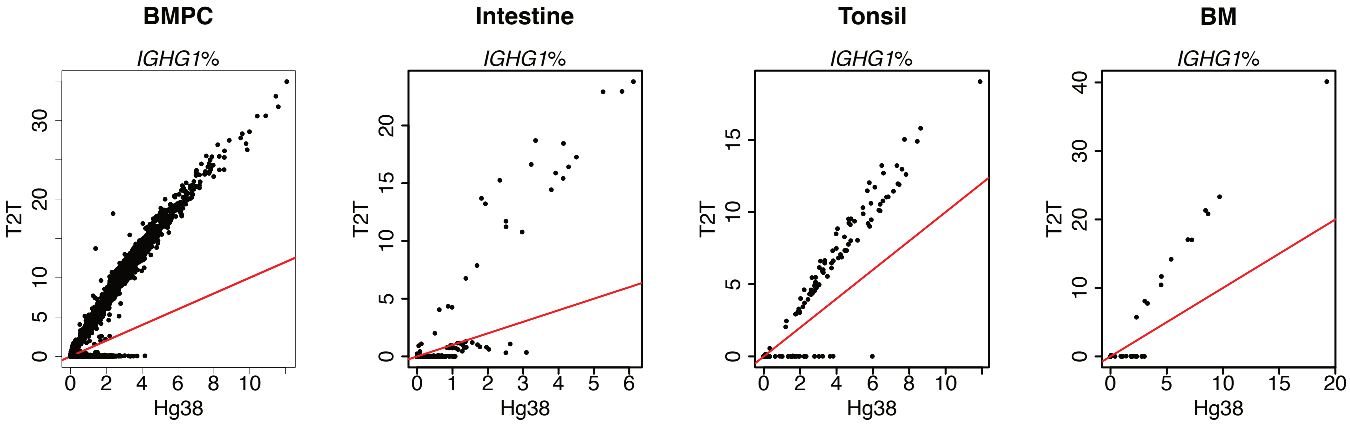


**Figure S15 The comparison of *IGHG1* between T2T and hg38**

**A)** Detailed alignment results for the CH3-CHS exons of *IGHG1* for both T2T and hg38. Mismatched bases are marked in red. 3-UTR regions are represented in lowercase. The green bar indicates the read of Fig S11C and the light-blue bar indicates the read of Fig 4E.

**B)** Percentage of *IGHG1* reads per cell for the BMPC, intestine, tonsil and bone marrow datasets. Hg38 and T2T results are compared by scatter plots. Red lines in scatter plots represent the x=y line.

### Figure S16

**A**

```
                    *         *         *         *         *         *         *         *         *         *
IGHG2-T2T    1>GGCAGCCCCGAGAACCACAGGTGTACACCCTGCCCCCATCCCGGGAGGAGATGACCAAGAACCAGGTCAGCCTGACCTGCCTGGTCAAAGGCTTCTACCC>100
IGHG2-hg38   1>GGCAGCCCCGAGAACCACAGGTGTACACCCTGCCCCCATCCCGGGAGGAGATGACCAAGAACCAGGTCAGCCTGACCTGCCTGGTCAAAGGCTTCTACCC>100

                    *         *         *         *         *         *         *         *         *         *
           101>CAGCGACATCGCCGTGGAGTGGGAGAGCAATGGGCAGCCGGAGAACAACTACAAGACCACACCTCCCATGCTGGACTCCGACGGCTCCTTCTTCCTCTAC>200
           101>CAGCGACATCTCCGTGGAGTGGGAGAGCAATGGGCAGCCGGAGAACAACTACAAGACCACACCTCCCATGCTGGACTCCGACGGCTCCTTCTTCCTCTAC>200

                    *         *         *         *         *         *         *         *         *         *
           201>AGCAAGCTCACCGTGGACAAGAGCAGGTGGCAGCAGGGGAACGTCTTCTCATGCTCCGTGATGCATGAGGCTCTGCACAACCACTACACACAGAAGAGCC>300
           201>AGCAAGCTCACCGTGGACAAGAGCAGGTGGCAGCAGGGGAACGTCTTCTCATGCTCCGTGATGCATGAGGCTCTGCACAACCACTACACACAGAAGAGCC>300

                    *         *         *         *         *         *         *         *         *         *
           301>TCTCCCTGTCTCCGGGTAAATGAgtgccacggccagcaagcccccgctccccaggctctcggggtcgcgcgaggatgcttggcacgtaccccgtgtacat>400
           301>TCTCCCTGTCTCCGGGTAAATGAgtgccacggccggcaagcccccgctccccaggctctcggggtcgcgcgaggatgcttggcacgtaccccgtctacat>400

                    *         *         *         *         *
           401>acttcccgggcacccagcatggaaataaagcacccagcgcttccctgggcccctgcga>458
           401>acttcccgggcacccagcatggaaataaagcacccagcgctgccctgggcccctgcga>458
```

**B**

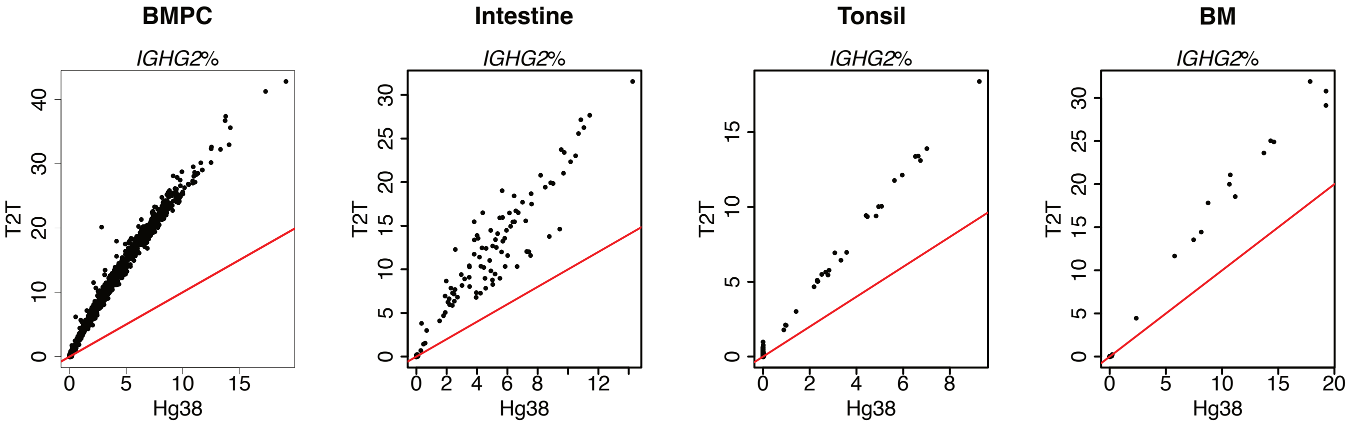


**Figure S16 The comparison of *IGHG2* between T2T and hg38**

**A)** Detailed alignment results for the CH3-CHS exons of *IGHG2* for both T2T and hg38. Mismatched bases are marked in red. 3-UTR regions are represented in lowercase. The green bar indicates the read of Fig 3C and the light-blue bar indicates the read of Fig 4F.

**B)** Percentage of *IGHG2* reads per cell for the BMPC, intestine, tonsil and bone marrow datasets. Hg38 and T2T results are compared by scatter plots. Red lines in scatter plots represent the x=y line.

### Figure S17

**A**

```
                          *         *         *         *         *         *         *         *         *         *
IGHG4-hg38     1>GGCAGCCCCGAGAGCCACAGGTGTACACCCTGCCCCCATCCCAGGAGGAGATGACCAAGAACCAGGTCAGCCTGACCTGCCTGGTCAAAGGCTTCTACCC>100
IGHG4-1-T2T    1>GGCAGCCCCGAGAGCCACAGGTGTACACCCTGCCCCCATCCCAGGAGGAGATGACCAAGAACCAGGTCAGCCTGACCTGCCTGGTCAAAGGCTTCTACCC>100
IGHG4-2-T2T    1>GGCAGCCCCGAGAGCCACAGGTGTACACCCTGCCCCCATCCCAGGAGGAGATGACCAAGAACCAGGTCAGCCTGACCTGCCTGGTCAAAGGCTTCTACCC>100

                          *         *         *         *         *         *         *         *         *         *
             101>CAGCGACATCGCCGTGGAGTGGGAGAGCAATGGGCAGCCGGAGAACAACTACAAGACCACGCCTCCCGTGCTGGACTCCGACGGCTCCTTCTTCCTCTAC>200
             101>CAGCGACATCGCCGTGGAGTGGGAGAGCAATGGGCAGCCGGAGAACAACTACAAGACCACGCCTCCCGTGCTGGACTCCGACGGCTCCTTCTTCCTCTAC>200
             101>CAGCGACATCGCCGTGGAGTGGGAGAGCAATGGGCAGCCGGAGAACAACTACAAGACCACGCCTCCCGTGCTGGACTCCGACGGCTCCTTCTTCCTCTAC>200

                          *         *         *         *         *         *         *         *         *         *
             201>AGCAGGCTCACCGTGGACAAGAGCAGGTGGCAGGAGGGGAATGTCTTCTCATGCTCCGTGATGCATGAGGCTCTGCACAACCACTACACACAGAAGAGCC>300
             201>AGCAGGCTAACCGTGGACAAGAGCAGGTGGCAGGAGGGGAATGTCTTCTCATGCTCCGTGATGCATGAGGCTCTGCACAACCACTACACGCAGAAGAGCC>300
             201>AGCAAGCTCACCGTGGACAAGAGCAGGTGGCAGGAGGGGAATGTCTTCTCATGCTCCGTGATGCATGAGGCTCTGCACAACCACTACACACAGAAGAGCC>300

                          *         *         *         *         *         *         *         *         *         *
             301>TCTCCCTGTCTCTGGGTAAATGAgtgccagggccggcaagcccccgctccccgggctctcggggtcgcgcgaggatgcttggcacgtaccccgtgtacat>400
             301>TCTCCCTGTCTCTGGGTAAATGAgtgccagggccggcaagcccccgctccccgggctctcggggtcgcgcgaggatgcttggcacgtaccccgtgtacat>400
             301>TCTCCCTGTCTCTGGGTAAATGAgtgccagggccggcaagcccccgctccccgggctctcggggtcgcgcgaggatgcttggcacgtaccccgtctacat>400

                          *         *         *         *         *
             401>acttcccgggcgcccagcatggaaataaagcacccagcgctgccctgggcccctgcga>458
             401>acttcccgggcgcccagcatggaaataaagcacccagcgctgccctgggcccctgcga>458
             401>acttcccaggcacccagcatggaaataaagcacccaccactgccctgggcccctgtga>458
```

**B**

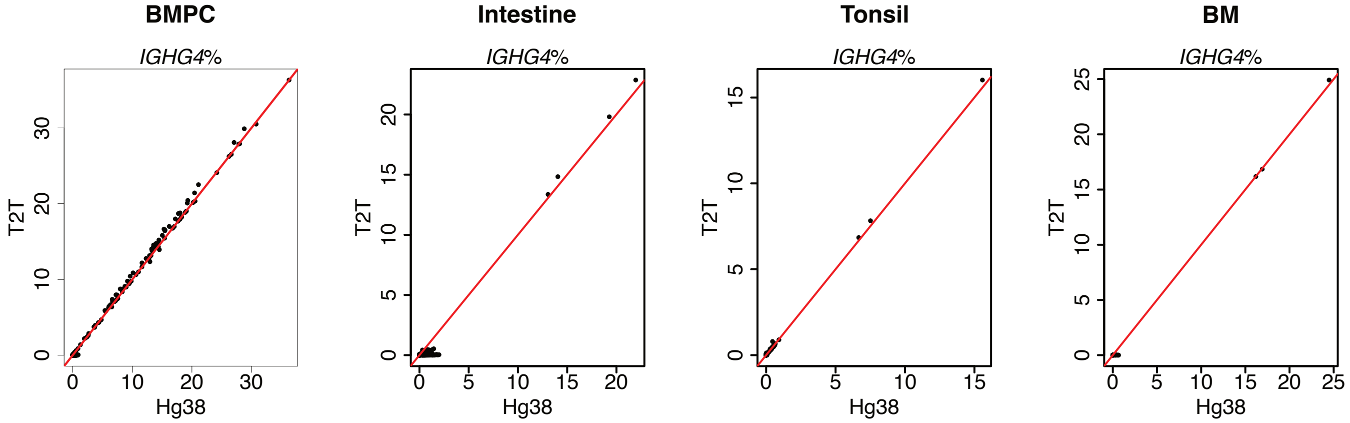


**Figure S17 The comparison of *IGHG4* between T2T and hg38**

**A)** Detailed alignment results for the CH3-CHS exons of *IGHG4* for both T2T (including two copies) and hg38. Mismatched bases are marked in red. 3-UTR regions are represented in lowercase.

**B)** Percentage of *IGHG4* reads per cell for the BMPC, intestine, tonsil and bone marrow datasets. Hg38 and T2T results are compared by scatter plots. Red lines in scatter plots represent the x=y line.

### Figure S18

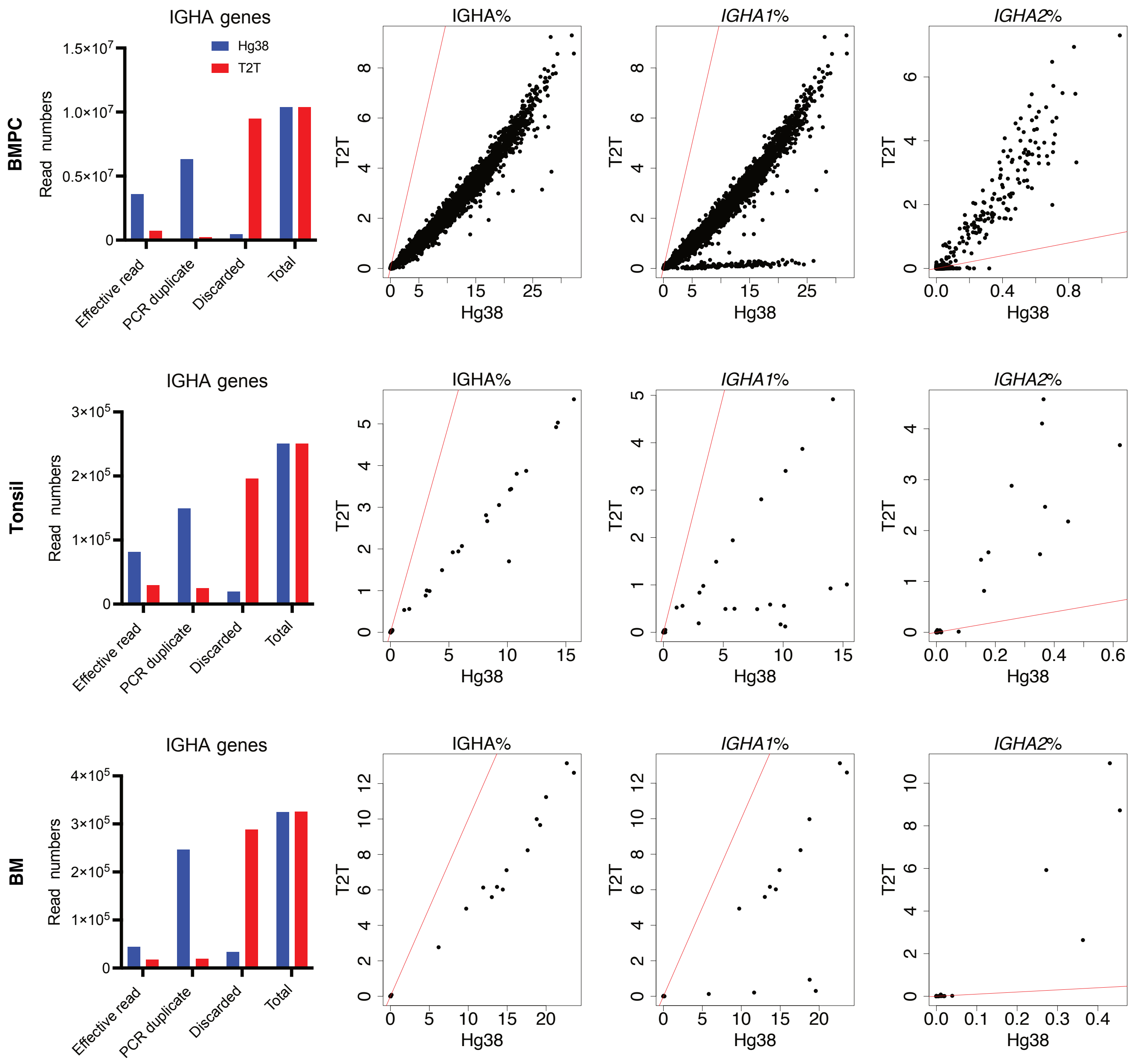


**Figure S18 T2T has fewer IGHA reads but better distinguishes *IGHA1* from *IGHA2***

The left barplots: Classification of reads that are assigned to IGHA genes for the BMPC (top), tonsil (middle) and bone marrow (bottom) datasets for each reference genome. The barplots show the total read count of IGHA genes and also the number of effective reads, the number of reads identified as PCR duplicates and the number of reads discarded for other reasons.

The right scatter plots: Percentage of IGHA genes reads per cell (left), percentage of *IGHA1* (middle) and *IGHA2* (right) reads per cell for the BMPC (top), tonsil (middle) and bone marrow (bottom) datasets. Hg38 and T2T results are compared by scatter plots. Red lines in scatter plots represent the x=y line.

### Figure S19

```
                      *         *         *         *         *         *         *         *         *         *
IGHA1-T2T    1>GAAACACATTCCGGCCCGAGGTCCACCTGCTGCCGCCGCCGTCGGAGGAGCTGGCCCTGAACGAGCTGGTGACGCTGACGTGCCTGGCACGCGGCTTCAG>100
IGHA2-T2T    1>GAAACACATTCCGGCCCGAGGTCCACCTGCTGCCGCCGCCGTCGGAGGAGCTGGCCCTGAACGAGCTGGTGACGCTGACGTGCCTGGCACGTGGCTTCAG>100
IGHA1-hg38   1>GAAACACATTCCGGCCCGAGGTCCACCTGCTGCCGCCGCCGTCGGAGGAGCTGGCCCTGAACGAGCTGGTGACGCTGACGTGCCTGGCACGCGGCTTCAG>100
IGHA2-hg38   1>GAAACACATTCCGGCCCGAGGTCCACCTGCTGCCGCCGCCGTCGGAGGAGCTGGCCCTGAACGAGCTGGTGACGCTGACGTGCCTGGCACGTGGCTTCAG>100

                      *         *         *         *         *         *         *         *         *         *
           101>CCCCAAGGACGTGCTGGTTCGCTGGCTGCAGGGGTCACAGGAGCTGCCCCGCGAGAAGTACCTGACTTGGGCATCCCGGCAGGAGCCCAGCCAGGGCACC>200
           101>CCCCAAGGATGTGCTGGTTCGCTGGCTGCAGGGGTCACAGGAGCTGCCCCGCGAGAAGTACCTGACTTGGGCATCCCGGCAGGAGCCCAGCCAGGGCACC>200
           101>CCCCAAGGATGTGCTGGTTCGCTGGCTGCAGGGGTCACAGGAGCTGCCCCGCGAGAAGTACCTGACTTGGGCATCCCGGCAGGAGCCCAGCCAGGGCACC>200
           101>CCCCAAGGATGTGCTGGTTCGCTGGCTGCAGGGGTCACAGGAGCTGCCCCGCGAGAAGTACCTGACTTGGGCATCCCGGCAGGAGCCCAGCCAGGGCACC>200

                      *         *         *         *         *         *         *         *         *         *
           201>ACCACCTTCGCTGTGACCAGCATACTGCGCGTGGCAGCCGAGGACTGGAAGAAGGGGGACACCTTCTCCTGCATGGTGGGCCACGAGGCCCTGCCGCTGG>300
           201>ACCACCTTCGCTGTGACCAGCATACTGCGCGTGGCAGCCGAGGACTGGAAGAAGGGGGACACCTTCTCCTGCATGGTGGGCCACGAGGCCCTGCCGCTGG>300
           201>ACCACCTTCGCTGTGACCAGCATACTGCGCGTGGCAGCCGAGGACTGGAAGAAGGGGGACACCTTCTCCTGCATGGTGGGCCACGAGGCCCTGCCGCTGG>300
           201>ACCACCTACGCTGTAACCAGCATACTGCGCGTGGCAGCTGAGGACTGGAAGAAGGGGGAGACCTTCTCCTGCATGGTGGGCCACGAGGCCCTGCCGCTGG>300

                      *         *         *         *         *         *         *         *         *
           301>CCTTCACACAGAAGACCATCGACCGCTTGGCGGGTAAACCCACCCATGTCAATGTGTCTGTTGTCATGGCGGAGGTGGACGGCACCTGCTACTGAgc-cg>399
           301>CCTTCACACAGAAGACCATCGACCGCTTGGCGGGTAAACCCACCCATGTCAATGTGTCTGTTGTCATGGCGGAGGTGGACGGCACCTGCTACTGAgc-cg>399
           301>CCTTCACACAGAAGACCATCGACCGCTTGGCGGGTAAACCCACCCATGTCAATGTGTCTGTTGTCATGGCGGAGGTGGACGGCACCTGCTACTGAgc-cg>399
           301>CCTTCACACAGAAGACCATCGACCGCATGGCGGGTAAACCCACCCACATCAATGTGTCTGTTGTCATGGCGGAGGCGGATGGCACCTGCTACTGAgcttg>400

               *         *          *         *               *
           400>cccgcctgtccccacccct-gaataaactcca-tgct-c-c-c-c-caagcag~>445
           400>cccgcctgtccccacccct-gaataaactcca-tgct-c-c-c-c-caagc~~~>443
           400>cccgcctgtccccacccct-gaataaactcca-tgct-c-c-c-c-caagcag~>445
           401>-g-g-g-g-a-gca----tggagttta-ttcaggggtggggacaggcgggcggc>443
```

**Figure S19 The sequence comparison of IGHA genes**

Detailed alignment results for the CH3-CHS exons of IGHA genes for both T2T and hg38. The red, orange, and blue bars refer to the corresponding regions of Fig 6G. The light-blue bar refers to *IGHA* exons of Fig 7A. Mismatched bases are marked in red. 3-UTR regions are represented in lowercase.

### Figure S20

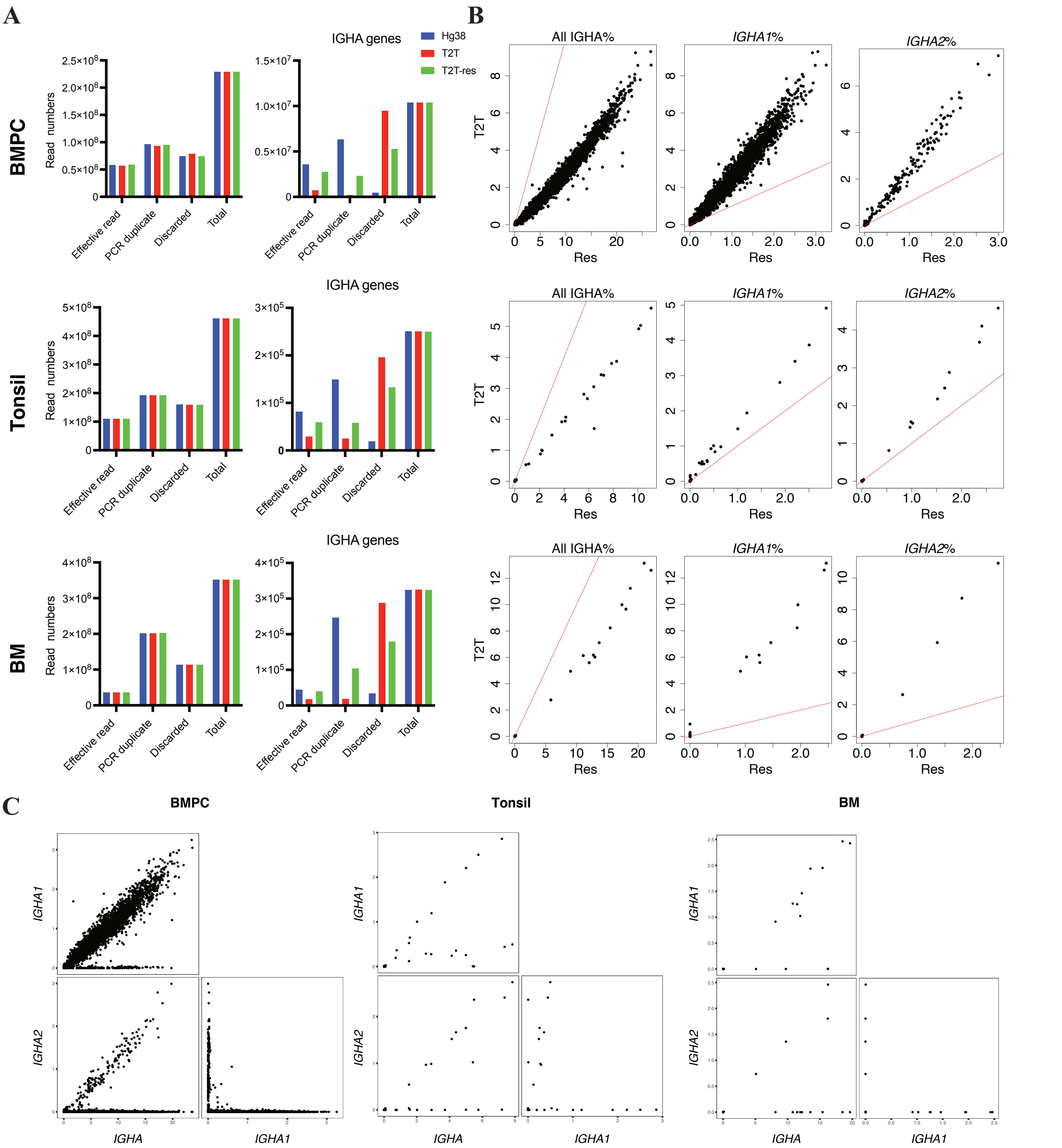


**Figure S20 A custom annotation reference for T2T can rescue multi-mapped reads mapped to IGHA genes**

**A)** Classification of reads (left) and reads that are assigned to IGHA genes (right) for the BMPC (top), tonsil (middle) and bone marrow (bottom) datasets for each reference. The barplots show the total read count for each sample and also the number of effective reads, the number of reads identified as PCR duplicates and the number of reads discarded for other reasons. Only effective reads were used for downstream analyses.

**B**) Percentage of IGHA genes reads per cell (left), *IGHA1* reads per cell (middle), and *IGHA2* reads per cell (right) for the BMPC (top), tonsil (middle) and bone marrow (bottom) datasets. T2T-res (shown as "res" in plots) and T2T results are compared by scatter plots. Red lines in scatter plots represent the x=y line.

**C)** Paired scatter plots correlating the expression levels of pairs of IgA subclass genes for the BMPC (left), tonsil (middle) and bone marrow (right) datasets. Plots are shown for the output using the custom reference based on T2T. Here expression is represented as reads per hundred (RPH).

### Figure S21

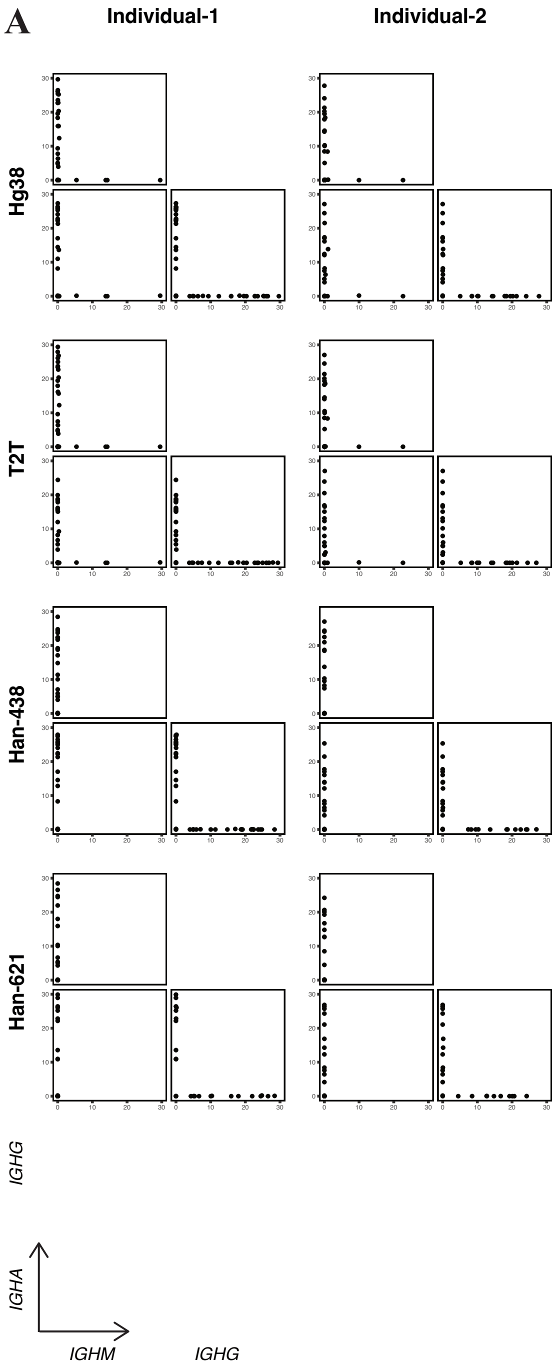


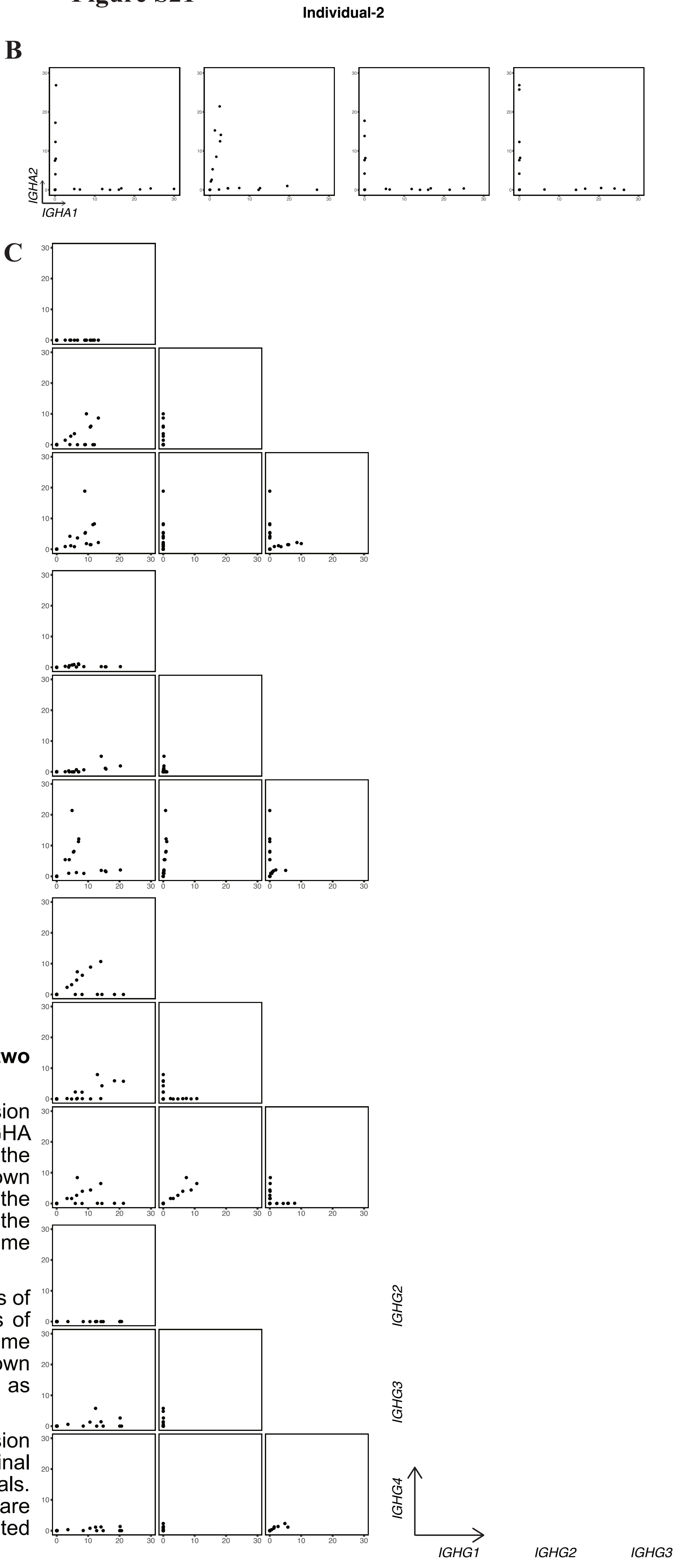


**Figure S21 The isotype gene expression of two Chinese individuals**

**A)** Paired scatter plots correlating the expression levels of pairs of *IGHM*, total IGHG, and total IGHA for the 2 Chinese individuals. The individual and the genome reference used of each plot is shown accordingly. There is no *IGHM* and *IGHD* in the annotations of Han-438 and Han-621, thus the expression level of *IGHM* is 0 for these two genome references.

**B)** Scatter plots correlating the expression levels of *IGHA1* and *IGHA2* for intestinal mucosa ASCs of the second Chinese individual. The genome references used of each plot are shown accordingly. Here expression is represented as reads per hundred.

**C)** Paired scatter plots correlating the expression levels of pairs of IgG subclass genes for intestinal mucosa ASCs of the second Chinese individuals. The genome references used of each plot are shown accordingly. Here expression is represented as reads per hundred.

**Table S1**: The numbers of gene x cell for each step of quality control as described in methods for the 4 datasets.

| | BMPC | | Intestine | | Tonsil | | BM | |
|---|---|---|---|---|---|---|---|---|
| | Hg38 | T2T | Hg38 | T2T | Hg38 | T2T | Hg38 | T2T |
| Raw | 20300x7459 | 20454x7469 | 20300x10200 | 20454x10007 | 20300x7460 | 20454x7457 | 20300x5996 | 20454x6000 |
| Step1 | 20300x7431 | 20454x7441 | 20300x9162 | 20454x8925 | 20300x7054 | 20454x7052 | 20300x5811 | 20454x5814 |
| Step2 | 20120x7431 | 20274x7441 | 20120x9162 | 20274x8925 | 20120x7054 | 20274x7052 | 20120x5811 | 20274x5814 |
| Step3 | 11371x7431 | 11354x7441 | 13341x9162 | 13311x8925 | 13588x7054 | 13585x7052 | 12661x5811 | 12671x5814 |
| Step4 | 11371x6790 | 11354x6785 | 13341x7905 | 13311x7813 | 13588x6601 | 13585x6602 | 12661x4815 | 12671x4818 |
| ASCs | 11371x6760 | 11354x6751 | 13341x1247 | 13311x1157 | 13588x209 | 13585x211 | 12661x46 | 12671x46 |
| Common ASCs | 6738 | | 1153 | | 209 | | 46 | |

### Computer code used to create Cell Ranger references, counts and mapping scores

##### 1. Generate FASTQ files for Cell Ranger

From SRR files:

```
fastq-dump --split-files --gzip -O fastq/ -v $srr
```

From BAM files:

```
${path_to_Cell_Ranger}/cellranger-7.0.0/lib/bin/bamtofastq --nthreads=$core $bam fastq/
```

##### 2. Make Cell Ranger reference

```
cellranger mkref --genome=${name} --fasta=$fa --genes=$gtf_edited --nthreads=$core --memgb=$mem
```

##### 3. Cell Ranger counts

```
cellranger count --id $id --transcriptome ${name} --fastqs fastq/ --localcores $core --expect-cells $num_cell --nosecondary
```

##### 4. Get read classification (xf), mapping scores (AS) and edit distance (nM)

```
samtools view ${id}/possorted_genome_bam.bam -@ $core | awk  -v tag=${tag} 'BEGIN {OFS=","} {for(i=12;i<=NF;++i) {if($i ~ "^" tag ":") {split($i, arr, ":"); print $1,arr[3]}}}' | gzip > output/${id}_${tag}.txt.gz

samtools view ${id}/possorted_genome_bam.bam -@ $core -D GN:gene_name.txt | awk  -v tag=${tag} 'BEGIN {OFS=","} {for(i=12;i<=NF;++i) {if($i ~ "^" tag ":") {split($i, arr, ":"); print $1,arr[3]}}}' | gzip > output/${id}_${tag}.txt.gz
```

##### 5. Create BAM files, get mapping scores and edit distances for double-positive cells

For Figure 3, Figure 5, Figure 10, and Figure 11 track:

```
samtools view $bam_hg38 -D CB:${cell_DP} -o hg38_cell.bam -@ $core

samtools view hg38_cell.bam -d GN:${gene1} -@ $core -b | samtools view -D xf:xf_tag.txt -@ $core -o hg38_cell_${gene1}.bam -

samtools index -@ $core hg38_cell_${gene1}.bam

samtools view hg38_cell.bam -d GN:${gene2} -@ $core -b | samtools view -D xf:xf_tag.txt -@ $core -o hg38_cell_${gene2}.bam -

samtools index -@ $core hg38_cell_${gene2}.bam

samtools view hg38_cell_${gene1}.bam -@ $core | cut -f1 > hg38_cell_${gene1}_readName.txt

samtools view hg38_cell_${gene2}.bam -@ $core | cut -f1 > hg38_cell_${gene2}_readName.txt

samtools view $bam_t2t -D CB:${cell} -o t2t_cell.bam -@ $core

samtools view t2t_cell.bam -N hg38_cell_${gene1}_readName.txt -D xf:xf_tag.txt -@ $core -o t2t_cell_hg38_cell_${gene1}.bam

samtools index -@ $core t2t_cell_hg38_cell_${gene1}.bam

samtools view t2t_cell.bam -N hg38_cell_${gene2}_readName.txt -D xf:xf_tag.txt -@ $core -o t2t_cell_hg38_cell_${gene2}.bam
```

```
samtools index -@ $core t2t_cell_hg38_cell_${gene2}.bam

samtools view -b -D xf:xf_tag.txt t2t_cell.bam > t2t_cell_xf.bam

samtools index t2t_cell_xf.bam
```

For Fig 4 and Fig 5 track:

```
samtools merge -o t2t_cell_hg38_cell_${gene1}_${gene2}.bam
t2t_cell_hg38_cell_${gene1}.bam t2t_cell_hg38_cell_${gene2}.bam

samtools index t2t_cell_hg38_cell_${gene1}_${gene2}.bam

samtools view t2t_cell_hg38_cell_${gene1}_${gene2}.bam | cut -f1 >
t2t_cell_hg38_cell_${gene1}_${gene2}_readName.txt

samtools view t2t_cell_xf.bam -d GN:${gene2} | cut -f1 >
t2t_cell_xf_${gene2}_readName.txt

Rscript - <<RSCRIPT

print("Running R")
a <- read.delim("t2t_cell_hg38_cell_${gene1}_${gene2}_readName.txt",header = F)
b <- read.delim("t2t_cell_xf_${gene2}_readName.txt",header = F)
c <- b[,1][!(b[,1] %in% a[,1])]
write(c,"t2t_cell_xf_${gene2}_extra_readName.txt")

RSCRIPT

samtools view -N t2t_cell_xf_${gene2}_extra_readName.txt t2t_cell.bam -o
t2t_cell_xf_${gene2}_extra.bam

samtools index t2t_cell_xf_${gene2}_extra.bam
```

Mapping scores and edit distances:

```
samtools view hg38_cell_${gene1}.bam | awk -v tag=${tag} 'BEGIN {OFS=","}
{for(i=12;i<=NF;++i) {if($i ~ "^" tag ":") {split($i, arr, ":"); print
$1,arr[3]}}}'

samtools view t2t_cell_hg38_cell_${gene1}.bam | awk -v tag=${tag} 'BEGIN {OFS=","}
{for(i=12;i<=NF;++i) {if($i ~ "^" tag ":") {split($i, arr, ":"); print
$1,arr[3]}}}'

samtools view hg38_cell_${gene2}.bam | awk -v tag=${tag} 'BEGIN {OFS=","}
{for(i=12;i<=NF;++i) {if($i ~ "^" tag ":") {split($i, arr, ":"); print
$1,arr[3]}}}'

samtools view t2t_cell_hg38_cell_${gene2}.bam | awk -v tag=${tag} 'BEGIN {OFS=","}
{for(i=12;i<=NF;++i) {if($i ~ "^" tag ":") {split($i, arr, ":"); print
$1,arr[3]}}}'
```

### R code used for Seurat analyses

#### 1. Read Cell Ranger counts and create Seurat object

```
library(Seurat)

count <- Read10X("filtered_feature_bc_matrix/")

sce <- CreateSeuratObject(counts = count)
```

#### 2. Quality control

```
sce[["percent.Ig"]] <- PercentageFeatureSet(sce, pattern =
"^IG[HKL][VDJCMDGAE]([1-9]|$)")
sce[["percent.ribo"]] <- PercentageFeatureSet(sce, pattern = "^M?RP[LS]")
sce[["percent.mito"]] <- PercentageFeatureSet(sce, pattern = "^MT-")

mitothres <- 10
genecellthres <- 0.005
genenumthres <- 450
countnumthres <- 1500
Igthres <- 10

sce <- subset(sce,subset= percent.mito<mitothres)

rib <- grep("^M?RP[LS]",rownames(sce))
nonrib <- rownames(sce)[-rib]
sce <- subset(x = sce,features=nonrib)

keep_by_gene <- which(rowSums(sce@assays$RNA@counts > 0) >=
ncol(sce@assays$RNA@counts)*genecellthres)

sce <- subset(x = sce,features=rownames(sce)[keep_by_gene])

sce <- subset(x = sce, subset= nFeature_RNA>=genenumthres)

sce <- subset(x = sce, subset= nCount_RNA>=countnumthres)

sce[["percent.Ig"]] <- PercentageFeatureSet(sce, pattern =
"^IG[HKL][VDJCMDGAE]([1-9]|$)")

sce <- subset(sce,percent.Ig>=Igthres)
```

When comparing genomes:

```
intersectcell <- intersect(colnames(sce_hg38),colnames(sce_T2T))
sce <- subset(sce,cells = intersectcell)
```

#### 3. t-SNE and UMAP plots

```
IGHC <- c("IGHG1","IGHG2","IGHG3","IGHG4","IGHA1","IGHA2","IGHE","IGHM","IGHD")
sce <- NormalizeData(sce)
sce <- FindVariableFeatures(sce, selection.method = "vst", nfeatures = 500)
VariableFeatures(object = sce)<-VariableFeatures(object = sce)[-
c(grep("^IG[HLK][VDJCADEGM]",VariableFeatures(sce)),grep("^TR[ABGD][VDJC]",Variabl
eFeatures(sce)))]
VariableFeatures(sce) <- c(VariableFeatures(sce),IGHC)
all.genes <- rownames(sce)
sce <- ScaleData(sce, features = all.genes,do.center = FALSE,do.scale = FALSE)
sce <- RunPCA(sce, features = VariableFeatures(object = sce))
dim.use <- 1:15
sce <- RunTSNE(sce, dims = dim.use, seed.use = 8888)
sce <- RunUMAP(sce, dims = dim.use, seed.use = 8888)
DimPlot(sce,reduction = "tsne")
DimPlot(sce,reduction = "umap")
```
